# Morphological, molecular, and functional evidence for a CNS-like oral nerve ring in the sea anemone *Nematostella vectensis*

**DOI:** 10.64898/2026.01.28.701697

**Authors:** Ruohan Zhong, Chris W. Seidel, Anna M.L. Klompen, Matthew C. Gibson

## Abstract

The emergence of centralized nervous systems reflects a major inflection point in evolution, enabling animals to integrate diverse inputs and coordinate complex behaviors. Neural centralization is typically associated with Bilateria, whereas their sister group, Cnidaria (jellyfish, anemones, and corals), has long been thought to rely on diffuse nerve nets mediating simple reflexes. This view, reinforced by limited anatomical and molecular data, has left unresolved whether cnidarians can form localized centers for neural processing, a question sharpened by the growing recognition of their diverse behavioral repertoires. Here we show that the sea anemone *Nematostella vectensis* possesses an oral nerve ring composed of ganglion-like condensations, a hallmark of centralized organization. These neurons are enriched for excitatory, inhibitory, and modulatory receptors but lack sensory or ciliary markers, yielding a molecular profile most consistent with bilaterian interneurons. Genetic disruption of a conserved inhibitory receptor subunit predominantly expressed in the oral nerve ring delayed the initiation of swallowing in a novel feeding paradigm, demonstrating a potential role in behavioral regulation. Together, these findings provide converging anatomical, molecular, and functional evidence that cnidarians can assemble localized integrative centers, suggesting that key elements of neural centralization predated the cnidarian–bilaterian split.

## Introduction

Animals coordinate behavior by concentrating neurons into localized integrative centers, a strategy exemplified by the bilaterian central nervous system (CNS). In contrast, cnidarians, the sister group to bilaterians, are traditionally portrayed as relying on diffuse nerve nets that mediate simple reflexes^1–4^ (Fig. 1a). This dichotomy has long framed centralization as a bilaterian innovation and left unresolved whether cnidarians can assemble structures that support local integrative processing.

**Fig. 1.**
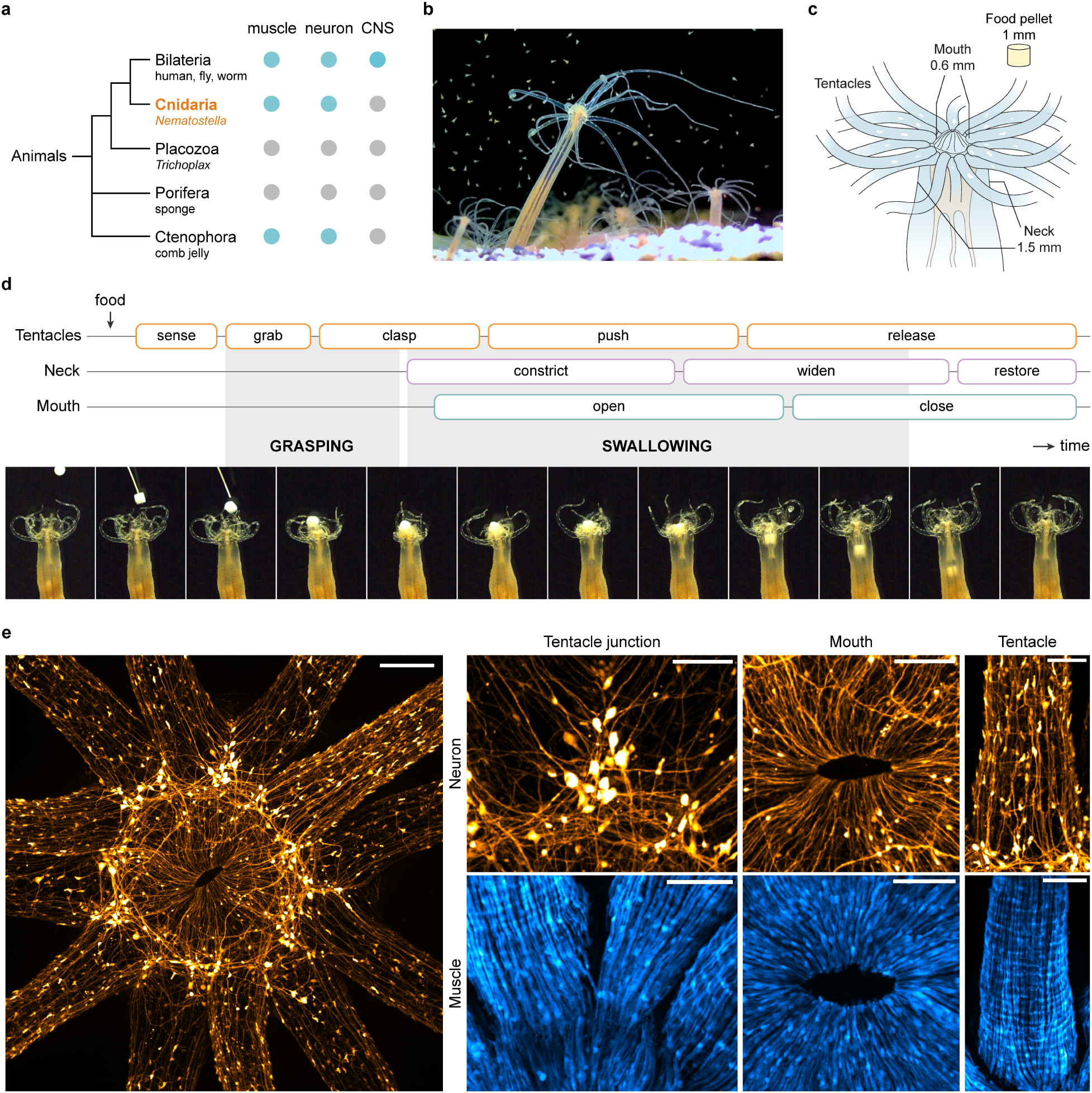
Structured motor coordination and a ganglion-like oral nerve ring in *Nematostella vectensis*. **(a)** Phylogenetic distribution of muscles, neurons, and centralized nervous systems (CNS) across major animal lineages; the dotted branch indicates unresolved phylogenetic relationships. **(b)** Adult *Nematostella vectensis* partially buried in gravel while capturing live brine shrimp (*Artemia*). A corresponding video is provided in Supplementary Video 1. **(c)** Schematic of the standardized feeding paradigm, showing the relative geometry of the oral disc, tentacles, and neck, and the size of the food pellet (1 mm). **(d)** Time-lapse montage illustrating two sequential phases of feeding, grasping and swallowing, separated by the onset of neck constriction. Each phase is composed of coordinated motor subroutines executed by the tentacles, neck, and mouth. A corresponding time-lapse video is provided in Supplementary Video 6. **(e)** Live confocal images of juvenile *Nematostella vectensis* expressing the neuronal reporter *NvSH3GL3^P^>mScarlet-I* (orange) or a muscle reporter driven by the myosin heavy chain promoter^51^ (blue). Neuronal and muscle images were acquired from separate animals and are shown for anatomical reference. Left, overview of the oral region showing dense neuronal labeling in the oral disc and tentacles. Right, enlarged views of selected regions reveal distinct neuronal morphologies and spatial organization. Neurons with enlarged somata relative to surrounding neurons cluster at tentacle junctions and extend circumferential neurites, forming a ring-like pattern. Additional neurons exhibit smaller somata and distinct projection orientations, with radial projections around the oral opening and longitudinal projections within tentacles. Scale bars, 100 µm (overview) and 50 µm (enlarged views).

The diffuse nerve net paradigm has persisted for over a century, in part due to assumptions tied to cnidarians’ basal phylogenetic position, the absence of macroscopic brain- or nerve cord-like structures, and a lack of molecular tools to visualize and classify neurons. *Hydra*, long the principal model for cnidarian neurobiology, further shaped this view through its diffuse, highly regenerative but lineage-specific neural organization^5,6^. Within this framework, cnidarian nerve nets were widely interpreted as morphologically and functionally simple. Still, classical anatomical studies hint that this picture might be incomplete. In 1879, Oscar and Richard Hertwig described a high density of neural elements arranged in crisscross and radial patterns in the oral disc of the parasitic anemone *Calliactis*^7^. In 1963, Elaine Robson illustrated radial oral nerve tracts in the swimming anemone *Stomphia* and proposed that they mediate coordination between the oral disc and the mesenteries^8^. In 1982, Kinnamon and Westfall identified interneuron-like cells at hypostome–tentacle junctions in *Hydra littoralis*, which they referred to as “primitive ganglia”^9^. These findings raise the possibility of localized neural organization within cnidarian nerve nets, although their cellular identities, developmental origins, and functional roles remain unknown.

Modern behavioral work strengthens this perspective by revealing functional specialization across cnidarian lineages. *Clytia hemisphaerica* relies on spatially organized subnetworks to coordinate food handling^10^; *Hydra vulgaris* displays satiety-dependent behaviors mediated by enteric- and CNS-like neuronal subpopulations^11^; the box jellyfish *Tripedalia cystophora* can modify obstacle avoidance through operant conditioning^12^; and the sea anemone *Nematostella vectensis* exhibits circadian locomotion and associative learning^13,14^. Together, these findings indicate that cnidarian nerve nets support more structured and context-dependent processing than is captured by the classical diffuse nerve net paradigm.

*Nematostella vectensis* offers an ideal system to revisit these assumptions with modern tools. It combines a tractable life cycle^15,16^, a well-annotated genome^17–19^, and an expanding molecular toolkit^20–22^. Unlike bilaterian invertebrates such as *Drosophila* or *C. elegans*, which have undergone extensive gene loss, *Nematostella* retains a broad ancestral eumetazoan gene repertoire more comparable to vertebrates, including synaptic machinery, neurotransmitter pathways, and developmental regulators^23–30^. Despite its compact genome (269.4 Mb; ∼19,200 protein-coding genes, similar to humans), our analysis of the current RefSeq assembly reveals an unexpectedly broad diversification of neurotransmitter receptor families, including glutamate, γ-aminobutyric acid (GABA), acetylcholine, monoamine, and neuropeptide receptors, well beyond the two previously characterized types^31,32^ and far exceeding the diversity seen in many bilaterians (Supplementary Table 2). In contrast, receptor numbers for other signaling pathways such as BMP^33^, Wnt^34^, and Hedgehog^35^ are comparable to or below human levels, suggesting that this diversification is specific to neural communication genes. Developmentally, *Nematostella* exhibits bilateral patterning of tentacles, musculature, and Hox/BMP expression along its directive axis^22,33,36^, undermining the traditional association between radial body plans and diffuse neural organization.

Whether cnidarians can assemble discrete interneuron-like centers capable of local integrative processing has remained unclear. Here, using anatomical, molecular, and functional approaches, we examine nervous system organization in the cnidarian *Nematostella vectensis* and assess whether features associated with neural centralization extend deeper in animal evolution than previously appreciated.

## Results

### Motor coordination during feeding

Feeding, an innate, essential, and readily inducible behavior requiring whole-body coordination, provides a tractable entry point to test whether a cnidarian nerve net contains localized neural structures that support complex motor transitions. To characterize feeding under ecologically relevant conditions, we recorded adult *Nematostella* polyps partially burrowed in gravel with the oral disc exposed above the substrate (Fig. 1b). Upon introduction of free-swimming *Artemia* nauplii (brine shrimp larvae), tentacles that made contact bent, twisted, and captured prey, whereas non-contacting tentacles remained poised in the background. Prey-bearing tentacles then contracted toward the oral opening, conveying food through the pharynx and into the body cavity (Supplementary Video 1).

In initial experiments, we found that feeding responses were highly flexible and context dependent, varying with prey size, number, position, and mode of delivery (Supplementary Videos 2–5 and Supplementary Fig. 5f). To identify the core elements within this behavioral diversity, we established a simplified feeding paradigm in which a cylindrical egg-white pellet (1 mm × 1 mm) was gently delivered to the oral disc using a cat whisker (Fig. 1c,d and Supplementary Video 6). This consistently triggered a reproducible sequence of motor subroutines involving distinct body regions. We defined two stereotyped phases: (1) grasping, from the first tentacle-driven movement to the onset of neck constriction, during which tentacles sense and ensnare the pellet, pull it toward the mouth, and clasp around it; and (2) swallowing, from the onset of neck constriction until the pellet reaches the distal end of the pharynx, characterized by rhythmic neck contractions, mouth opening and closing, food passage through the pharynx, and temporary suspension of oral-to-aboral peristalsis. Our observations thus demonstrate that *Nematostella* executes a structured, multi-step feeding program despite lacking any obvious anatomical centralization.

### Anatomical organization of the oral nerve net

The adaptability and robustness of feeding under varied conditions raise the question of how coordinated behaviors can be directed by a diffuse nerve net. In *Nematostella*, a previously described *NvElav1* reporter labels neurons along the body column^25^ but shows little expression in neurons of the oral disc and tentacles. To overcome this limitation, we generated additional reporter lines by screening promoters of candidate genes identified in scRNA-seq data^28^.

Among them, a 1.9 kb upstream fragment of the *Nematostella SH3GL3* (Endophilin-A3) homolog drove a fluorescent reporter *NvSH3GL3^P^>mScarlet-I* (Fig. 2b) that robustly labeled neurons, including dense innervation of the oral disc and tentacles, major structures involved in feeding (Fig. 1e). Notably, this pattern resembles early hand-drawn descriptions of oral neuronal condensations in other sea anemones^7,8^, suggesting that it may represent a conserved anatomical feature.

**Fig. 2.**
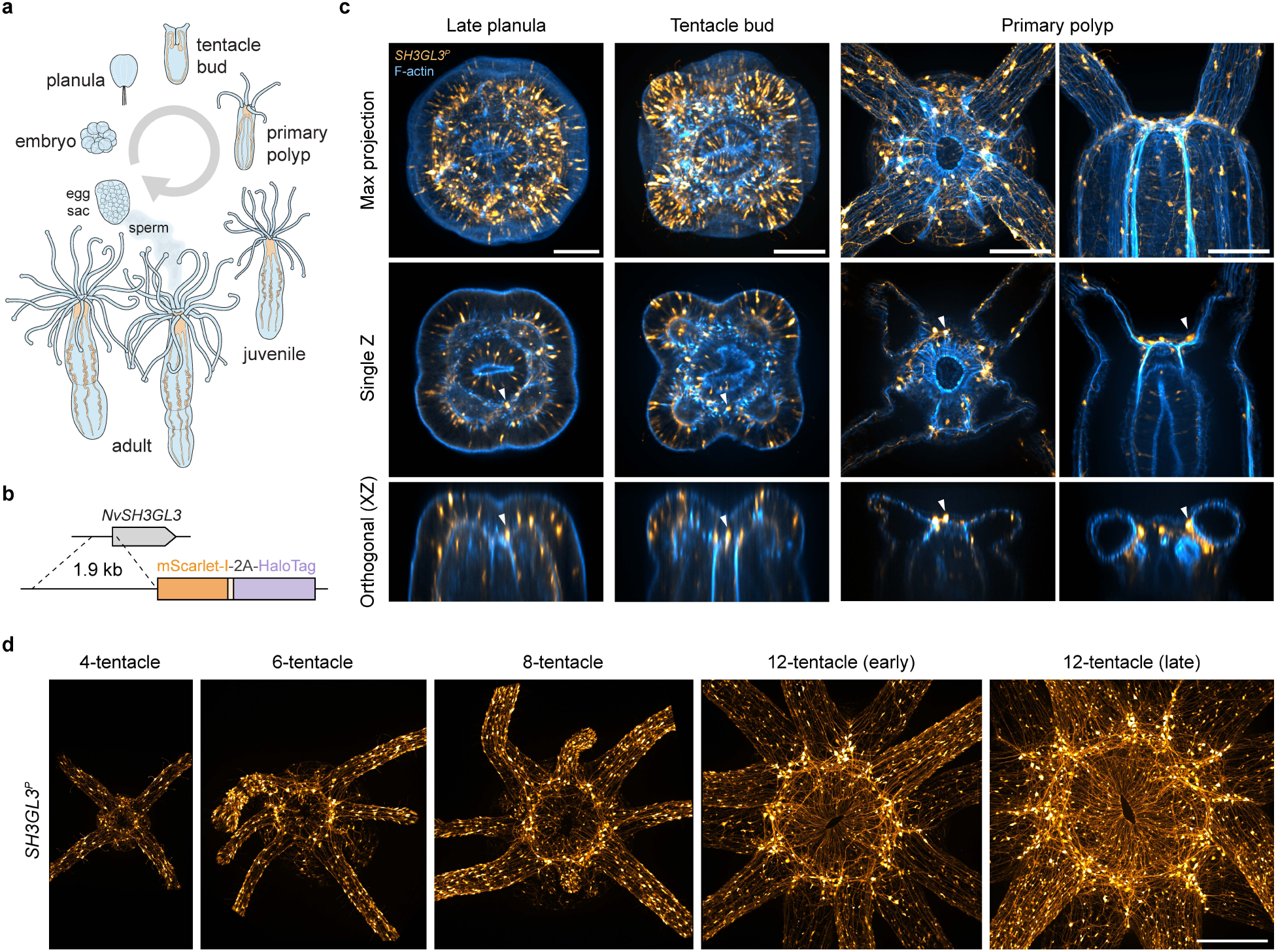
Developmental emergence and scaling of the oral nerve ring. **(a)** Schematic overview of *Nematostella vectensis* life stages, from embryo to adult, indicating the progressive addition of tentacles and the transition from planula larva to primary polyp and juvenile stages. **(b)** Schematic of the *NvSH3GL3^P^* reporter construct. A 1.9 kb upstream regulatory region of *NvSH3GL3* drives expression of an mScarlet-I fluorescent reporter used to visualize neuronal morphology. **(c)** Confocal images showing *NvSH3GL3^P^>mScarlet-I*–labeled neurons (orange) across early developmental stages, with F-actin staining (blue). Columns show late planula (3.5 days post-fertilization, dpf), tentacle buds (4.5 dpf), and primary polyps (10 dpf). Rows show maximum intensity projections (top), single optical sections (middle), and orthogonal (XZ) views (bottom). Late planula and tentacle-bud stages are shown in oral view, whereas primary polyps are shown in both oral and lateral views. In late planulae, sparsely distributed neurons with enlarged somata are observed in basal positions within the oral ectoderm. As tentacles emerge during the tentacle-bud and primary polyp stages, the number of these large-soma neurons increases, accompanied by higher neurite density and localization to the ectoderm at tentacle junctions. White arrowheads indicate representative neuronal somata shown in corresponding single optical sections and orthogonal views. Scale bars, 50 μm. **(d)** Oral-view maximum intensity projections showing *NvSH3GL3^P^>mScarlet-I*–labeled neurons (orange) in polyps with increasing tentacle number (4, 6, 8, and 12). As polyps grow and sequentially add tentacles, the circumorally projecting neurite bundle that constitutes the oral nerve ring (ONR) becomes progressively denser, accompanied by increased accumulation of large neuronal somata at tentacle junctions. All panels are shown at the same scale. Scale bar, 200 μm.

High-resolution imaging of the oral region revealed that neuronal organization is not diffuse but rather exhibits pronounced spatial patterning in both soma distribution and neurite orientation (Fig. 1e). Neurons with enlarged somata and multipolar morphology clustered at tentacle junctions, where they gave rise to prominent circumferential neurite bundles that together formed a ring-like structure. We refer to this neuronal structure as the oral nerve ring (ONR). In contrast, other oral neurons exhibited smaller somata and distinct projection orientations, including radially oriented neurites around the mouth and longitudinal projections within the tentacles (Fig. 1e). With circumferential neurites linking tentacles and localization at the interface between the tentacles, pharynx, and body column, the ONR occupies a position well suited for coordinating activity across multiple body regions.

To examine how this circumoral organization arises, we analyzed *NvSH3GL3^P^*-labeled neurons during larval-to-polyp transitions associated with tentacle formation (Fig. 2a,c). In late planula larvae, neurons with enlarged somata were present but sparsely distributed within the oral ectoderm and did not exhibit a coherent ring-like arrangement. As animals progressed through the tentacle-bud stage and into primary polyps, the number of these ONR neurons increased, accompanied by higher neurite density and preferential localization to tentacle junctions (Fig. 2c). With further growth and the sequential addition of tentacles^36^, this process gave rise to a progressively denser circumoral neurite bundle and an increased accumulation of large neuronal somata at tentacle junctions, ultimately forming a clearly defined oral nerve ring (Fig. 2d). These observations indicate that ONR organization emerges gradually during development and scales with increasing body size and morphological complexity.

### Molecular diversity of the oral nerve net

Building on the anatomical and developmental characterization of the oral nerve ring, we next examined its molecular composition to assess potential functional specialization. Oral discs from adult *NvSH3GL3^P^* reporter animals were dissected, and mScarlet-I-positive cells were enriched by fluorescence-activated cell sorting (FACS) prior to single-cell RNA sequencing using the 10x Genomics platform (Fig. 3a). This yielded 3,297 cells comprising both neural and non-neural populations (Fig. 3b), as defined by marker gene expression (Fig. 3c). Specifically, neural clusters expressed canonical neuronal markers (*ELAV*, *achaete-scute*), synaptic machinery (*synapsin*, *synaptotagmin*, *endophilin*, vesicular transporters), as well as cytoskeletal proteins, ion channels, transporters, and adhesion/signaling molecules (Fig. 3c and Supplementary Fig. 1a). Expression of the *NvSH3GL3^P^* reporter closely paralleled endogenous *NvSH3GL3* across neural clusters, supporting the fidelity of the reporter. In contrast, non-neural clusters expressed the endogenous red fluorescent protein *NvFP-7R*^37^ together with tissue-associated cytoskeletal and extracellular-matrix genes, providing an internal reference for non-neural identity (Fig. 3c).

**Fig. 3.**
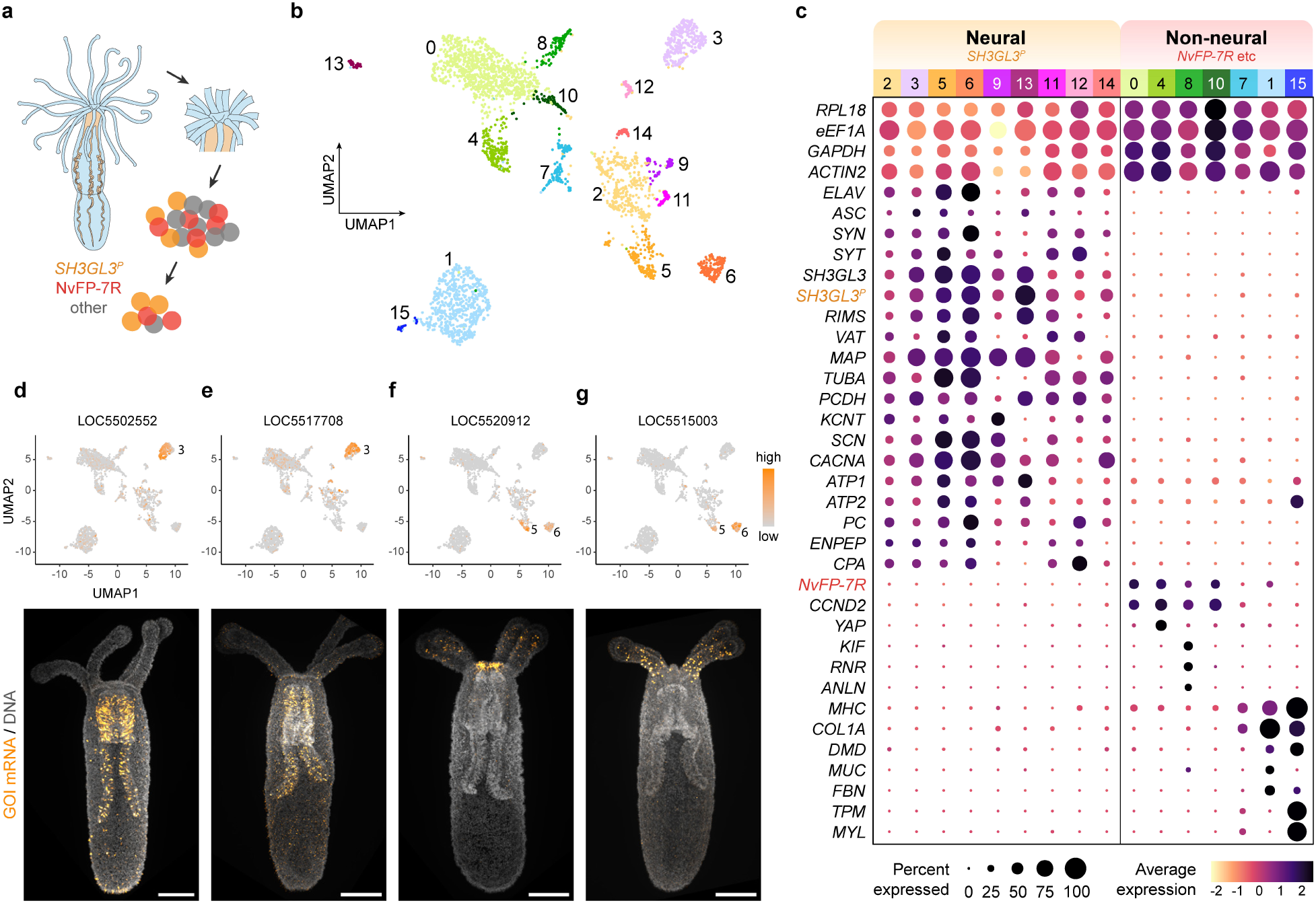
Molecular diversity and spatial organization of the oral nerve net. **(a)** Schematic of oral-end dissection and cell enrichment for single-cell RNA sequencing. Oral ends from adult *Nematostella vectensis* were dissected, and most tentacles were trimmed prior to mechanical and enzymatic dissociation for 10x Genomics single-cell RNA sequencing. Cells were enriched for *NvSH3GL3^P^>mScarlet-I*-positive populations by fluorescence-activated cell sorting (FACS), with gating applied to reduce spectral contamination from cells expressing the native red fluorescent protein NvFP-7R. Colors indicate major cell populations (orange, *NvSH3GL3^P^>mScarlet-I*; red, NvFP-7R; gray, other). **(b)** UMAP visualization of 3,297 cells from the single-cell dataset. **(c)** Dot plot showing representative gene categories, including housekeeping genes, canonical neuronal and synaptic markers, cytoskeletal components, ion channels and transporters, and adhesion/signaling molecules. Dot size indicates the percentage of cells expressing each gene, and color represents scaled average expression (pale yellow to dark purple). Neural clusters (left) show strong *SH3GL3^P^*expression, whereas non-neural clusters (right) include *NvFP-7R*-enriched populations as well as clusters enriched for contractile, cytoskeletal, and extracellular-matrix genes. Expression of the *SH3GL3^P^* reporter parallels endogenous *SH3GL3*, supporting reporter fidelity. A complete list of neural genes with identifiers is provided in Supplementary Fig. 1a. **(d–g)** Feature plots (top) show SCT-normalized gene expression projected onto the UMAP, with corresponding fluorescent in situ hybridization (FISH) images (bottom) in four-tentacle primary polyps (7–10 dpf) for four representative cluster-enriched genes. (d,e) LOC5502552 and LOC5517708 are enriched in cluster 3 and display predominant expression in the pharyngeal region by FISH. (f,g) LOC5520912 and LOC5515003 are expressed in partially overlapping subsets of clusters 5 and 6 and show predominant FISH signal in the oral disc and tentacles. Feature plots are scaled independently for each gene, with light gray indicating no detectable expression and increasing color intensity indicating higher expression. DNA is shown in gray, and gene-specific signal is shown in orange. Scale bars, 100 µm.

To examine the spatial expression of markers enriched in specific transcriptional clusters, we analyzed four-tentacle primary polyps (7–10 days post-fertilization, dpf), which are more amenable to whole-mount fluorescent in situ hybridization (FISH) and confocal imaging than adult tissues. Markers enriched in cluster 3 consistently localized to the pharyngeal region (Fig. 3d,e). In contrast, markers enriched in clusters 5 and 6 localized predominantly to the ectoderm of the oral disc and tentacles (Fig. 3f,g and Supplementary Fig. 1b–e). Together, these findings show that at least a subset of transcriptionally defined neural clusters within the oral nerve net exhibit clear spatial correlates across oral tissues, linking molecular identity to anatomical position.

### Interneuron-like molecular features of the oral nerve ring

We next examined the molecular diversity of oral neurons, as reflected in their sensory transduction components and neurotransmitter receptor repertoires, to assess whether transcriptionally defined populations exhibit features consistent with functional specialization. Sensory-related genes included acid-sensing ion channels (ASICs), transient receptor potential (TRP) channels, opsins, and cilia-associated proteins (CFAPs, DNAHs, and RSPs). Neurotransmitter receptors fell into three classes: excitatory (ionotropic glutamate, acetylcholine), inhibitory (ionotropic GABA, glycine), and modulatory (metabotropic glutamate, acetylcholine, GABA, monoamines, neuropeptides). Full gene lists are provided in Supplementary Table 2.

Comparison across transcriptional clusters revealed that clusters 5 and 6 were distinguished by low sensory gene expression and broad representation of excitatory (E), inhibitory (I), and modulatory (M) receptor classes (Fig. 4a). Co-expression analysis showed that the majority of cells within these clusters simultaneously expressed receptors from all three classes (Fig. 4b and Supplementary Fig. 2), indicating a molecular profile consistent with integrative signal processing rather than sensory specialization, reminiscent of integrative interneurons in the bilaterian CNS. This molecular profile also mirrored two morphological features observed in ONR neurons labeled by the *NvSH3GL3^P^*reporter, including the absence of sensory cilia and the presence of ganglion-like neuronal condensations. Although clusters 5 and 6 shared these general characteristics, their receptor compositions diverged. Cluster 5 exhibited substantial heterogeneity in E/I/M receptor combinations, whereas cluster 6 displayed a more uniform and balanced receptor profile across cells (Fig. 4c). This balance, approximately 20–30–50 in the relative proportions of excitatory, inhibitory, and modulatory receptor expression per cell, was maintained despite variation in receptor identity, suggesting functional convergence within cluster 6.

**Fig. 4.**
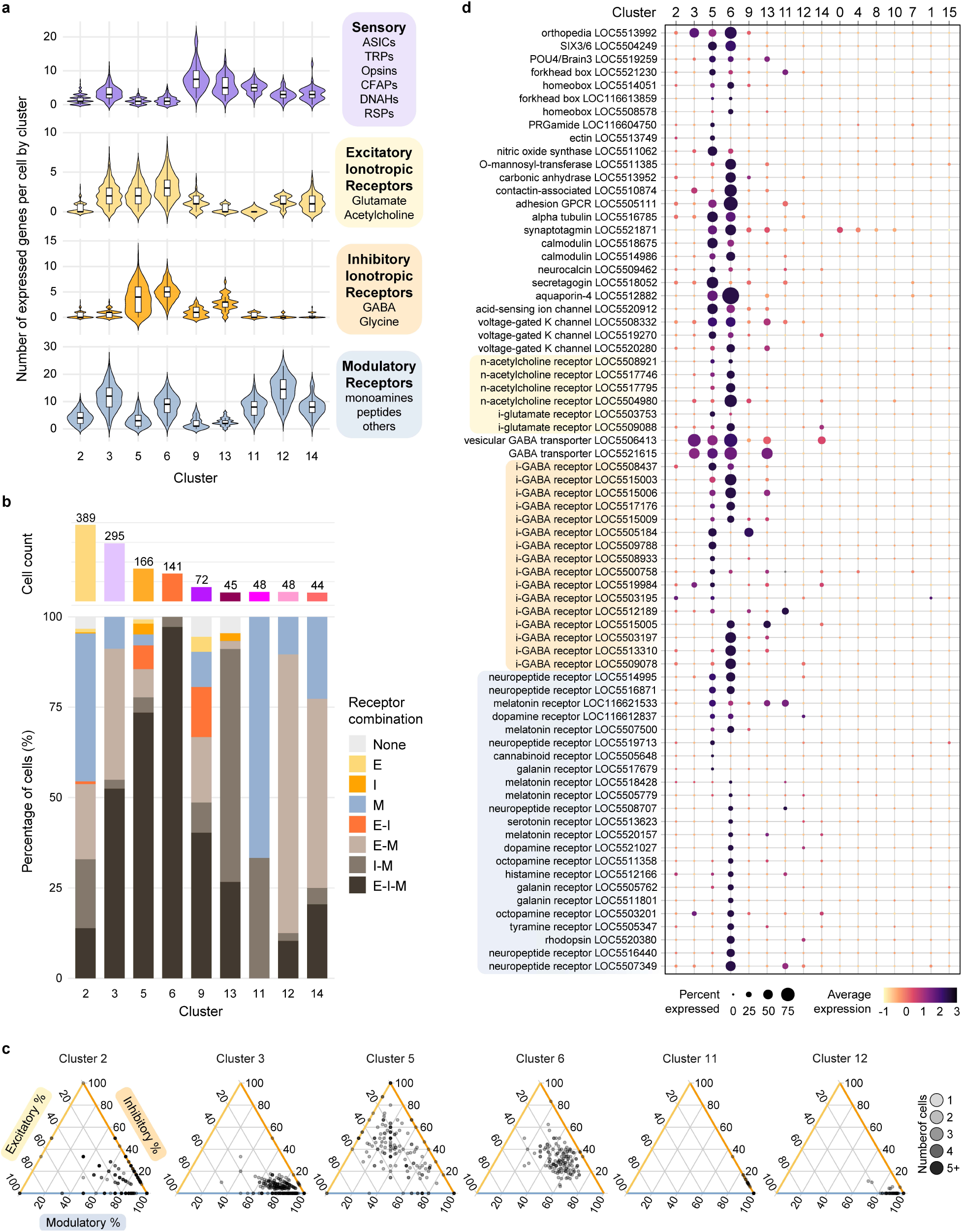
Integrative receptor repertoires distinguish interneuron-like neural clusters. **(a)** Violin plots showing the number of sensory-related and neurotransmitter receptor genes detected per cell across neural clusters. Gene categories are color-coded as sensory (purple), ionotropic excitatory (E, gold), ionotropic inhibitory (I, orange), and metabotropic/modulatory (M, blue). **(b)** Co-expression analysis of E/I/M receptor combinations across neural clusters. The upper histogram shows total cell counts per cluster, and the lower stacked bar plot shows the percentage of cells in each cluster belonging to different receptor-class combination categories. Pale gray denotes cells lacking detectable neurotransmitter receptors; gold, excitatory-only; orange, inhibitory-only; blue, modulatory-only; orange-red, excitatory–inhibitory; light gray, excitatory–modulatory; medium gray, inhibitory–modulatory; and dark gray, cells co-expressing all three receptor classes. **(c)** Ternary plots showing the relative proportions of detected E, I, and M receptors per cell for selected clusters. Each point represents a cell, positioned according to its E/I/M receptor composition. Point color indicates the number of overlapping cells sharing the same receptor ratio, from light gray (1 cell) to black (≥ 5 cells). **(d)** Dot plot showing expression patterns of transcription factors, synaptic components, ion channels, and neurotransmitter receptors associated with fast neurotransmission and neuromodulation across clusters. Dot size indicates the percentage of cells expressing each gene, and color represents scaled average expression (pale yellow to dark purple). Gold, orange, and blue shading highlight genes belonging to the E, I, and M receptor families, respectively.

Other oral neural clusters differed from clusters 5 and 6 by showing sensory and/or modulatory biases rather than integrative receptor profiles. Cluster 9 exhibited a sensory bias, expressing receptors from all three classes, but with most cells single- or double-positive and enriched for sensory and ciliary genes (Fig. 4a,b). Clusters 11, 12, and 14 displayed a combined sensory and modulatory bias, characterized by distinct modulatory receptor expression at the cluster level (Supplementary Fig. 3a) and, at the single-cell level, by co-expression of broad sets of modulatory receptor genes together with sensory-related genes (Fig. 4a and Supplementary Fig. 3b–d). Despite these receptor-rich profiles, none of these clusters exhibited the balanced E/I/M co-expression pattern that distinguishes clusters 5 and 6 (Fig. 4a–c).

Marker analysis further reinforced these distinctions (Fig. 4d). Clusters 5 and 6 shared transcription factors including *POU4/Brn3*^29^, *Orthopedia (Otp)*, and *SIX3/6*^27^, which are known to regulate anterior CNS patterning and neuronal specification in Bilateria^38–40^. Although *SIX3/6* expression is restricted to the aboral pole in *Nematostella* larvae^27^, it was enriched in adult oral disc neurons (Supplementary Fig. 1b). Both clusters also expressed multiple ionotropic GABA receptor-like subunits, with shared and cluster-specific paralogs (Fig. 4d and Fig. 6a), indicating subtype-specific inhibitory tuning. By contrast, cluster 2 showed broad neuronal gene expression without defining markers, consistent with an undifferentiated state. Together, these comparisons identified clusters 5 and 6 as the most plausible ONR-associated neuronal populations within the broader oral neural domain, based on their spatial localization and interneuron-like molecular profiles.

To identify molecular markers within clusters 5 and 6 that label ONR neurons, we performed FISH on four-tentacle primary polyps for genes enriched in these clusters. Candidate genes were excluded from further consideration if they fell into three categories that limited their utility as ONR markers: (1) genes enriched in the adult oral disc scRNA-seq dataset but weak or undetectable in primary polyps (Supplementary Fig. 1b); (2) transcripts detected in ONR-like cells but also expressed broadly along the body column (Supplementary Fig. 1c,d); and (3) transcripts confined to the oral end but either diffusely expressed across tissues or localized to cells lacking the large somata and tentacle-junction positioning characteristic of ONR neurons (Supplementary Fig. 1e). By contrast, multiple ionotropic GABA receptor-like (GABA_A_R-like) subunits satisfied both transcriptional enrichment and anatomical localization criteria, with expression concentrated in the oral region and in cells with ONR-like morphology (Fig. 3g, Fig. 6b, and Supplementary Fig. 1d). Although no single gene exclusively marked the entire ONR, these results highlight GABA signaling as a promising entry point for molecular and functional interrogation of the ONR.

### GABAergic pathway in the oral nerve net

Building on this molecular signature, we asked whether the ONR contains the core components of GABAergic signaling. In bilaterians, canonical GABAergic signaling involves four key elements: synthesis by glutamate decarboxylase (GAD), vesicular packaging by vesicular GABA transporter (VGAT), reception through ionotropic or metabotropic GABA receptors, and clearance by GABA transporter (GAT)^41^. To assess whether *Nematostella* possesses a comparable pathway architecture, we combined immunohistochemistry, single-cell and bulk transcriptomics, and comparative genomics.

Whole-mount immunostaining of *NvSH3GL3^P^>mScarlet-I* primary polyps revealed widespread GABA immunoreactivity, with strong enrichment at the oral end (Fig. 5a). Within the oral disc, a subset of GABA-positive cells overlapped with mScarlet-I-positive neurons possessing large somata characteristic of ONR morphology (Fig. 5b), suggesting that ONR neurons may both release and respond to GABA.

**Fig. 5.**
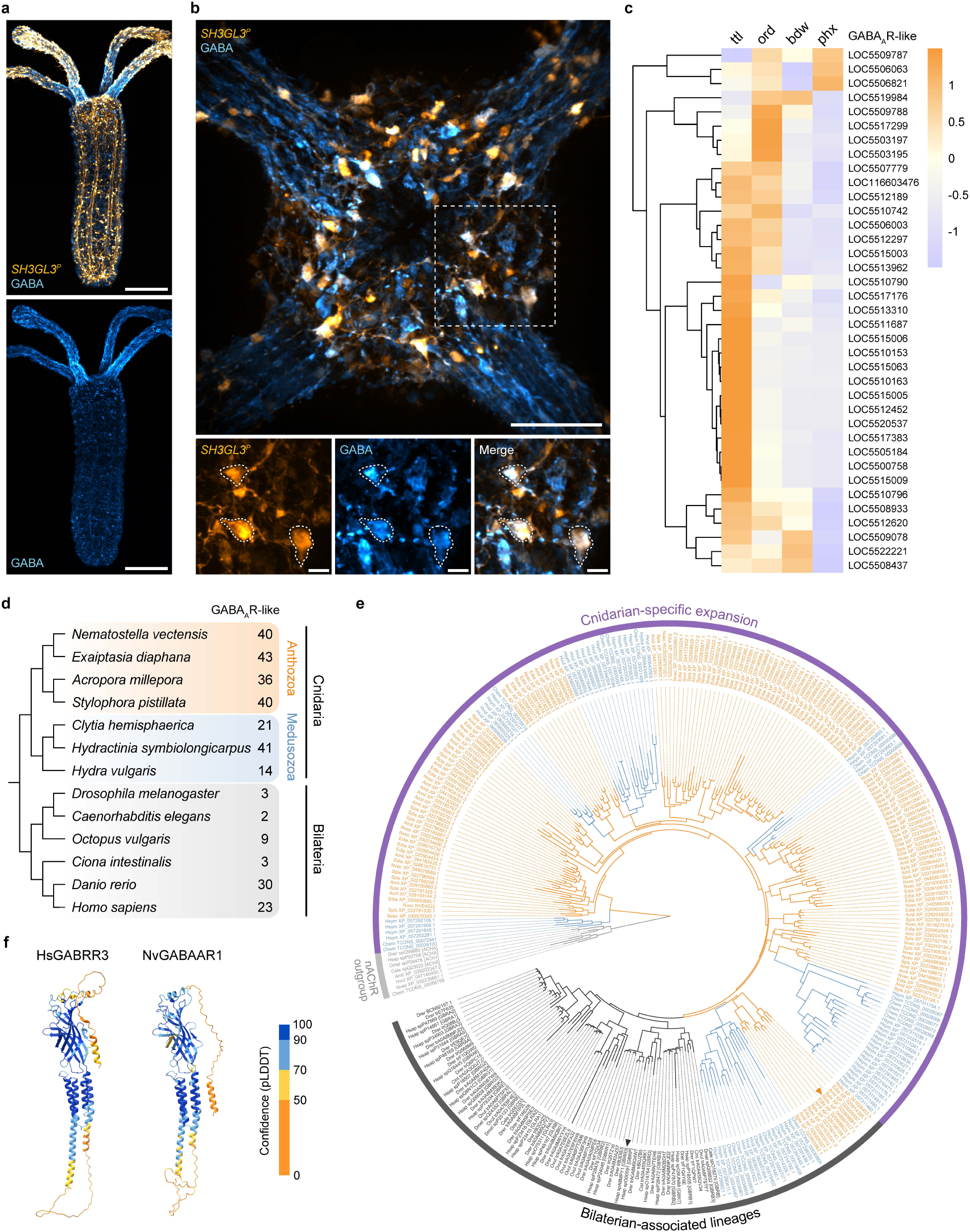
Expansion and oral enrichment of ionotropic GABAergic signaling components. **(a)** Whole-mount γ-aminobutyric acid (GABA) immunostaining of *NvSH3GL3^P^>mScarlet-I* primary polyps, showing enriched GABA immunoreactivity in the oral disc and tentacles. Scale bars, 100 µm. **(b)** Within the oral disc, GABA immunoreactivity is restricted to large-soma cells, including both mScarlet-I-positive neurons and additional large-soma GABA-positive cells lacking reporter signal. Top, merged image of *SH3GL3^P^* (orange) and GABA (blue), with the dashed box indicating the region shown below. Bottom, split-channel views of the boxed region (*SH3GL3^P^*, left; GABA, middle; merge, right). Dashed outlines mark representative large-soma cells that are double-positive for *SH3GL3^P^* and GABA. GABA accumulation is also visible in varicosities along neurites. Scale bars, 50 µm (top), 10 µm (bottom). **(c)** Heatmap showing tissue-specific expression of GABA type A receptor-like (GABA_A_R-like) homologs in adult *Nematostella*. Colors indicate Z-scored average TPM per gene across tissues (ttl, tentacles; ord, oral disc; bdw, oral body wall; phx, pharynx), with warmer colors representing higher relative expression. **(d)** Gene counts of GABA_A_R-like homologs across seven cnidarian species (four anthozoans, three medusozoans) and six bilaterian species. Anthozoans (orange) and medusozoans (blue) show marked expansion of the receptor family relative to bilaterians (dark gray). **(e)** Maximum-likelihood phylogeny of GABA_A_R-like homologs based on amino-acid sequences from the species set in (d). Anthozoan sequences are shown in orange, medusozoan sequences in blue, and bilaterian sequences in dark gray; representative nicotinic acetylcholine receptor (nAChR) subunits (light gray) form an outgroup to root the tree. Outer arcs denote the cnidarian-specific expansion (orange) and the bilaterian-associated lineage (dark gray). Arrowheads indicate the *Nematostella* subunit NvGABAAR1 (orange) and the human subunit GABRR3 (dark gray) selected for structural comparison in (f). Branch support for the cnidarian–bilaterian split was assessed with ultrafast bootstrap and SH-like aLRT values of 92.3 and 80; full support values are shown in Supplementary Fig. 4. **(f)** AlphaFold-predicted structures of *Homo sapiens* GABRR3 (γ-aminobutyric acid type A receptor subunit ρ3) and its *Nematostella vectensis* homolog NvGABAAR1 (protein ID: XP_048587017.1; gene ID: LOC5505184), corresponding to the subunits marked by arrowheads in (e). Ribbon coloring indicates per-residue model confidence (pLDDT): orange (0–50), yellow (50–70), light blue (70–90), and dark blue (90–100).

In our scRNA-seq dataset, among vesicular inhibitory amino acid transporter (VIAAT) homologs, only a single homolog showed substantial expression in adult oral neurons (LOC5506413; hereafter *VGAT*). Multiple GABA_A_R-like subunits showed expression largely confined to clusters 5 and 6 (Fig. 4d). Individual cells co-expressed 1–10 subunits (Fig. 4a), a combinatorial diversity rarely seen in invertebrates but characteristic of the vertebrate CNS^41,42^. Subunits displayed broad to cluster-specific distributions, with mutually exclusive, partially overlapping, or nested expression patterns (Fig. 6a), consistent with cell type-specific receptor assemblies. Notably, approximately 50% of cluster 5 neurons and 75% of cluster 6 neurons co-expressed GABA_A_R(s) and *VGAT* (Supplementary Fig. 5a), a defining feature of inhibitory interneurons capable of both releasing and responding to GABA^41^. Some GABA_A_R-positive neurons lacked *VGAT*, likely representing postsynaptic targets of inhibition. Together, these findings indicate that the ONR supports a GABAergic signaling architecture encompassing both presynaptic and postsynaptic components, with receptor diversity suggesting a capacity for fine-tuned inhibition comparable to bilaterian central circuits.

**Fig. 6.**
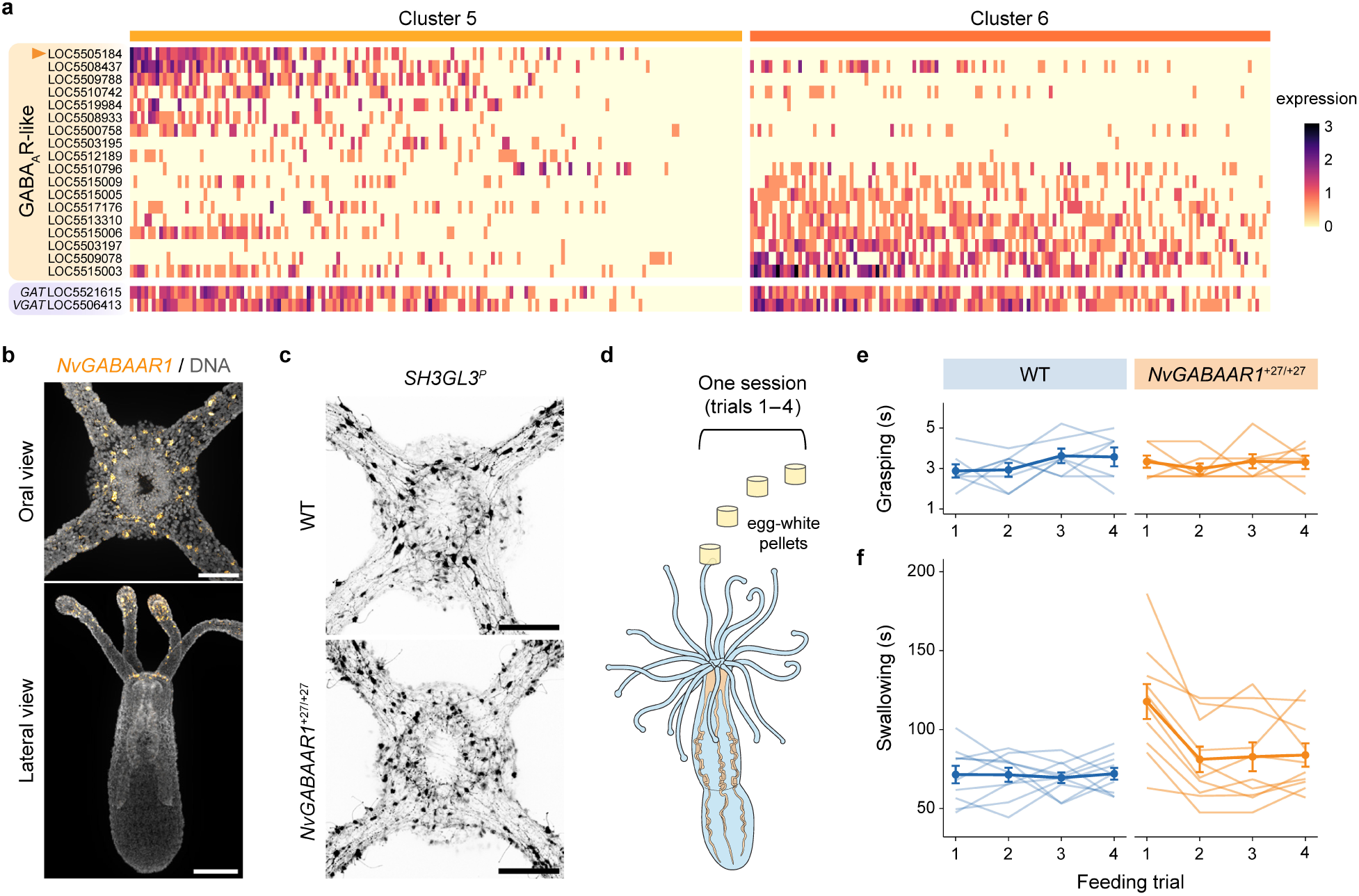
Loss of *NvGABAAR1* selectively delays swallowing initiation during feeding. **(a)** Heatmap showing SCT-normalized expression of genes involved in GABA signaling across single cells from clusters 5 and 6. The gene set includes multiple GABA_A_R-like subunits as well as the plasma membrane GABA transporter (*GAT*) and vesicular GABA transporter (*VGAT*). Columns represent individual cells ordered within each cluster by aggregate expression of this gene set. Rows are grouped by gene category and arranged from cluster 5-enriched (top) to cluster 6-enriched (bottom). The orange arrowhead marks LOC5505184 (*NvGABAAR1*), the subunit highlighted in Fig. 5e and analyzed functionally in Fig. 6b–f. **(b)** FISH in four-tentacle primary polyps showing *NvGABAAR1* expression restricted to the oral region, including the oral disc and a subset of tentacle neurons. *NvGABAAR1* signal in orange; DNA counterstain in gray. Top, oral view; bottom, lateral view. Scale bars, 50 µm (top), 100 µm (bottom). **(c)** Confocal images of wild-type (WT) and *NvGABAAR1*^+27/+27^ primary polyps in the *NvSH3GL3^P^>mScarlet-I* reporter background. The organization of the oral nerve ring appears comparable at the level of gross morphology between WT and mutants. Scale bars, 50 µm. **(d)** Feeding assay design. Each behavioral session consisted of four sequential presentations of 1 mm × 1 mm egg-white pellets delivered to the oral disc. **(e)** Grasping duration across four feeding trials using pipette-delivered food pellets. Thin lines represent individual animals (WT in blue, n = 7; *NvGABAAR1*^+27/+27^ mutants in orange, n = 7); bold lines and error bars denote mean ± s.e.m. Grasping duration was comparable between genotypes. **(f)** Swallowing duration across four feeding trials. Animals were assayed using either pipette or whisker delivery (each animal was tested with a single method throughout its session). Delivery method did not affect the genotype × trial interaction (see Methods), so data were analyzed together. Thin lines represent individual animals (WT in blue, n = 10; mutants in orange, n = 10); bold lines and error bars denote mean ± s.e.m. Mutants exhibited a pronounced increase in swallowing duration on trial 1 (46.6 ± 10.1 s; p = 0.0001) but converged to WT-like performance on trials 2–4. Full statistical details are provided in Supplementary Table 8.

To complement the single-cell RNA-seq data, we used bulk RNA-seq to transcriptionally profile the adult tentacles, oral disc, pharynx, and oral body wall. This approach provided deeper and less biased transcript coverage across whole tissues than scRNA-seq restricted to FACS-enriched oral neurons. GABA_A_R-like transcripts showed highest relative expression in the oral disc and tentacles, mirroring GABA immunostaining and reinforcing the receptor diversity of the oral nerve net (Fig. 5c).

Prompted by this enrichment, we surveyed the full *Nematostella* GABA_A_R-like gene set. Here, we use “GABA_A_R-like” to denote subunits with sequence similarity to bilaterian GABA_A_ or glycine receptor subunits, as detailed in the Methods. Genomic analysis identified 40 distinct subunits in *Nematostella*—exceeding the 2–23 typical of bilaterians (Fig. 5d and Supplementary Table 4). Phylogenetic analysis across 13 species (7 cnidarians and 6 bilaterians; Supplementary Table 5) supported a single ancestral GABA_A_R, with subsequent expansions in both lineages (Fig. 5e and Supplementary Fig. 4). In *Nematostella,* only three subunits clustered with bilaterian orthologs, whereas the remaining 37 arose from anthozoan-specific duplications, generating a lineage-specific expansion that underpinned ONR receptor diversity. A recent preprint reported a similar cnidarian-specific radiation based on narrower taxon sampling (2 cnidarians and 3 bilaterians) and functional assays of selected subunits^43^. Although metabotropic GABA receptors are also expanded (12 in *Nematostella* vs. 2 in humans; Supplementary Table 4), their expression was scattered across neural and non-neural clusters without ONR enrichment; therefore, we focused on ionotropic receptors as the core ONR-associated repertoire.

Despite the presence of both plasma membrane (GAT) and vesicular (VGAT/VIAAT) GABA transporters, and a diverse repertoire of GABA receptors, we did not identify a canonical *GAD* homolog enriched in neuronal populations within the oral scRNA-seq dataset. Ten putative *GAD*-like candidates, including two reported previously^44,45^, showed no enrichment in any neuronal cluster and were instead weakly or sparsely expressed in a small number of non-neural clusters (Supplementary Table 6 and Supplementary Notes). These findings suggest that GABA production in *Nematostella* may rely on non-canonical or lineage-divergent pathways.

### A GABA_A_ receptor-like subunit regulates swallowing initiation

The molecular profiling described above highlights GABA signaling as central to ONR function. Among the 40 GABA_A_R-like homologs encoded in the *Nematostella* genome, we focused on *NvGABAAR1* (LOC5505184), which showed the broadest expression within cluster 5 and the highest transcript levels (Fig. 6a). *NvGABAAR1* was also co-expressed with *VGAT*, a canonical feature of inhibitory interneurons. Moreover, it was one of only three GABA_A_R subunits that clustered with bilaterian orthologs (Fig. 5e), and AlphaFold modeling revealed strong structural similarity to human GABRR3 (Fig. 5f and Supplementary Fig. 5c). Crucially, FISH confirmed that *NvGABAAR1* was expressed in oral disc neurons with large somata and circumferential neurites, consistent with ONR morphology, although expression was also observed in a subset of tentacle neurons (Fig. 6b).

To test the function of *NvGABAAR1*, we used CRISPR/Cas9 mutagenesis to introduce a +27 bp insertion, generating a premature stop codon in the ligand-binding domain (Supplementary Fig. 5b). Homozygous *NvGABAAR1*^+27/+27^ animals were viable and displayed no overt neural or developmental defects (Fig. 6c).

We next assayed the behavior of *NvGABAAR1*^+27/+27^ mutants using our standardized feeding paradigm. Animals were sequentially presented with four sham prey items (1 mm × 1 mm egg-white pellets), and grasping and swallowing durations were quantified (Fig. 6d). Grasping duration, defined as the interval from the first tentacle-driven movement to the onset of neck constriction (Fig. 1d), did not differ between wild-type (WT) and mutant animals (p = 0.35), indicating intact prey detection and initial motor responses (Fig. 6e). Grasping duration was also stable across repeated trials for both genotypes, with no evidence of systematic drift over ten consecutive presentations (Supplementary Fig. 5d).

By contrast, swallowing showed a genotype-dependent effect restricted to the first of four feeding trials (Fig. 6f). Linear mixed-effects modeling revealed a significant genotype × trial interaction. Strikingly, WT animals showed stable swallowing durations across trials, whereas mutants exhibited a delay on the first feeding trial (46 ± 10 s; p < 0.001) and were indistinguishable from WT on subsequent trials (all p > 0.2). Importantly, including food-delivery method (pipette vs whisker) or starvation level as covariates did not alter the genotype × trial interaction, indicating that the delay on trial 1 reflects a robust, intrinsic deficit in swallowing initiation rather than an effect of stimulus delivery or feeding state. Mutant animals recovered rapidly after the initial trial and maintained WT-like swallowing durations for the remainder of the assay, a pattern that persisted for at least nine consecutive trials (Supplementary Fig. 5e). Together, these findings demonstrate that *NvGABAAR1* is specifically required for the initiation of swallowing behavior, supporting a role for GABAergic signaling within the oral nerve net in coordinating the transition from prey capture to ingestion and, potentially, the engagement of feeding-related behavioral states.

## Discussion

Cnidarian nerve nets have long been viewed as diffuse and reflexive, yet their capacity for structured, context-dependent behaviors hints at deeper organization. Here we show that the sea anemone *Nematostella vectensis* possesses an oral nerve ring (ONR), a localized neural structure implicated in motor coordination during feeding. This finding suggests that key elements of neural integration predate the cnidarian–bilaterian split some 700 million years ago, extending the evolutionary roots of centralized organization.

The ONR forms a localized aggregation of neuronal somata that meets classical criteria for a ganglion-like structure. In bilaterians, ganglia are clusters of somata functioning as local integrative or relay centers, whereas in cnidarians “ganglion cells” traditionally refer to aciliate neurons positioned basally within the epithelium^2,25^. Unlike previously described neurite condensations in cnidarian nerve nets^25,46,47^, the ONR constitutes a true ganglion-like aggregation of neuronal somata, providing the first modern evidence of soma-level clustering in a cnidarian (Fig. 1e). We use “ganglion-like” here in the morphological sense only, to describe neuronal clustering rather than positional homology to bilaterian ganglia. This clustered architecture provides an anatomical basis for a CNS-like site of integration within a diffuse nerve net. ONR neurons display morphological diversity, including bipolar and tripolar forms, suggesting heterogeneity in connectivity and function. Developmentally, the ONR arises from initially scattered neurons that progressively cluster as polyps grow (Fig. 2c,d). This scaling of organization with body size and muscular compartmentalization may reflect an adaptive route toward more efficient long-range coordination, a pattern that parallels proposed evolutionary transitions from diffuse to centralized networks across early metazoans^4^. While postsynaptic partners remain unmapped, the ONR’s position and organization indicate an interneuron-like hub mediating sensory–motor coordination.

Molecular and functional evidence support an interneuron-like identity for ONR neurons. Single-cell analyses show that ONR neurons lack sensory markers but express a combinatorial set of excitatory, inhibitory, and modulatory receptors together with transcription factors typical of bilaterian interneurons (Fig. 4a–d). This transcriptional profile defines a CNS-like integrative module, in which prominent inhibitory components, including GABA-immunoreactive somata, a vesicular GABA transporter, and an expanded suite of GABA_A_ receptor-like subunits, form a molecular framework for regulating excitation balance and temporal precision^41^. Although comprehensive functional mapping of the ONR remains beyond the scope of this study, our functional perturbation of an ONR-enriched GABA_A_R subunit selectively delayed the onset of swallowing behavior without impairing prey detection or grasping (Fig. 6e,f). The delay was most evident during the first feeding event but normalized by the second trial, indicating rapid behavioral recovery once feeding was initiated. This transient phenotype suggests that the ONR contributes to the timing and coordination of motor transitions during the initial stages of feeding rather than executing the motor output itself. Future work may uncover how internal physiological states shape ONR-mediated coordination. Together, these observations point to a role for inhibitory signaling within the ONR in modulating behavioral sequence progression, a hallmark of integrative rather than purely reflexive circuits.

Intriguingly, ONR neurons also expressed *SIX3/6* and *Orthopedia* (Fig. 4d), two broadly conserved transcription factors that in bilaterians regulate anterior CNS patterning and neuroendocrine specification^39,40^. These findings suggest that transcriptional programs defining the bilaterian anterior neuroectoderm may have deep evolutionary roots in the patterning of neural condensations around feeding structures.

Collectively, our results position the *Nematostella* oral nerve ring as an intermediate level of neural organization between diffuse nerve nets and centralized nervous systems. Its architecture exemplifies how clusters of interneuron-like cells can enable coordination without a bona fide brain. The marked expansion of neurotransmitter receptors (Supplementary Table 2) underscores molecular elaboration as a route to circuit flexibility in early animals, perhaps through receptor repurposing or adaptive tuning to estuarine variability. While it remains unresolved whether the *Nematostella* ONR and bilaterian CNS share a common origin or represent convergent solutions, their shared features reveal deep evolutionary continuity in the principles of neural integration.

## Methods

### Animals and husbandry

*Nematostella vectensis* anemones were maintained in 12 parts per thousand (ppt) artificial seawater (ASW, Instant Ocean) in glass containers at room temperature (RT, ∼22 °C) under low ambient light. Animals were fed freshly hatched *Artemia* nauplii 2–3 times per week, and seawater was replaced the following day to remove residual food and waste. Juvenile polyps were raised at 27 °C after their first feeding until sexual maturity, after which adults were kept long-term at either RT or 16 °C as needed. Mixed-sex populations were housed at the Stowers Institute and were originally derived from founder stock provided by Mark Martindale. Unless otherwise specified, all animals used in this study were kept under these standard conditions.

### Spawning and embryo preparation

Spawning was induced following established *Nematostella* protocols^48,49^. Sexually mature adults were kept at 16 °C, then transferred to RT and placed in front of a light box overnight to induce gamete release. The following morning, dishes were cold-shocked by replacing the water with 16–18 °C 12 ppt ASW. Unfertilized egg sacs were collected for microinjection. Egg sacs were dejellied in 4% L-cysteine prepared in 12 ppt ASW and incubated for 10 min on a gentle rocker. Wide-bore transfer pipettes were used to minimize mechanical damage. Dejellied eggs were washed several times in ASW, transferred to clean dishes, and kept at 16 °C until microinjection or fertilization.

Fertilized embryos used for fluorescent in situ hybridization and developmental imaging were cultured at 24 °C in the dark under standard conditions. Developmental stages used in this study included planula larvae (3 days post-fertilization, dpf) and primary polyps (7–10 dpf). Primary polyps corresponded to the first sessile feeding stage and were unfed, four-tentacle animals within this early post-metamorphic window. Juveniles were staged by tentacle number. Adults were considered as sexually mature polyps, typically bearing 16 tentacles and older than three months.

### Transgenic line generation

The *NvSH3GL3^P^>mScarlet-I* reporter construct was designed using a 1.9 kb promoter fragment upstream of the *NvSH3GL3* locus to drive an *mScarlet-I–CAAX–P2A–HaloTag* reporter cassette followed by an SV40 polyadenylation signal. mScarlet-I fluorescence was used for all anatomical imaging. The vector was assembled using the NEBuilder HiFi DNA Assembly Master Mix (NEB, E2621) and sequence-verified prior to injection. Oligonucleotides used for promoter cloning are listed in Supplementary Table 9.

Unfertilized oocytes were microinjected following established Meganuclease (I-SceI)-mediated transgenesis procedures^21,50^. Injection mix consisted of 10× I-SceI buffer (10 mM Tris-HCl, 10 mM MgCl_2_, 1 mM DTT, pH 8.8), ∼50 ng μl^-1^ plasmid DNA, 0.4 U μl^-1^ I-SceI enzyme (NEB, R0694), and FITC (Fisher Scientific, 46425) to visualize injection boluses. The mixture was assembled and pre-incubated according to standard protocol^21^. Microinjection was performed using an Eppendorf FemtoJet, and injected oocytes were fertilized after the injection session (≤ 1 h). Embryos were kept at 18 °C overnight and subsequently raised at RT with regular ASW changes until first feeding.

Injected animals retaining mosaic reporter expression after two feedings were pooled and raised to adulthood. Individual F0 adults were crossed to wild-type (WT) animals to assess germline transmission. A single F0 female carrier was identified and used to establish the line. Reporter-positive F1 offspring were outcrossed to WT animals, yielding ∼50% reporter-positive F2 progeny maintained as a stable heterozygous population. F2 reporter animals were used for all anatomical imaging and single-cell applications. F3 animals were generated by intercrossing F2 individuals, and homozygotes were identified by test-crossing to WT animals, which yielded 100% reporter-positive progeny. Homozygous F3 animals were maintained as genetic stock.

For anatomical reference in selected live-imaging experiments, a previously generated transgenic reporter expressing GFP under the control of the myosin heavy chain promoter^51^ was used to visualize muscle tissue. This reporter was used only for qualitative anatomical comparison and was not employed for quantitative analyses.

### CRISPR/Cas9 mutant generation

CRISPR/Cas9 genome editing was performed as previously described^20,52^. A single guide RNA (sgRNA) targeting the *NvGABAAR1* coding sequence was designed using the CRISPRscan server^53^ (https://www.crisprscan.org/sequence/) with default parameters (Cas9 - NGG, In vitro T7 promoter). The selected sgRNA targeted a site within the ligand-binding domain. A double-stranded DNA template for sgRNA synthesis was generated by annealing the gene-specific forward primer to a universal reverse primer, followed by Klenow extension (37 °C for 30 min, 75 °C for 15 min; Klenow Fragment, NEB, M0210). In vitro transcription was performed using the AmpliScribe T7-Flash Transcription Kit (Biosearch Technologies, ASF250) at 37 °C for 5 h, followed by DNase I treatment for 15 min. The sgRNA was purified using the Direct-zol RNA MiniPrep Kit (Zymo Research, R2052) according to the manufacturer’s instructions and quantified using a NanoDrop spectrophotometer (Thermo Fisher Scientific).

For microinjections, Cas9 protein with NLS (PNA Bio, CP02) was mixed with sgRNA at equal concentrations and diluted to 500 ng μl^-1^ each in nuclease-free water. FITC (Fisher Scientific, 46425) was added to visualize injection boluses. Unfertilized oocytes were injected with Cas9–sgRNA complexes, fertilized with WT sperm following the injection session, and raised at RT.

To evaluate editing efficiency and identify potential founders, a subset of injected embryos was sacrificed at 3–10 dpf for genotyping, since their small size precludes non-lethal tissue sampling. The remaining injected siblings were raised to adulthood as candidate F0 animals. A single F0 male was identified as carrying a +27 bp insertion, resulting in a premature stop codon within the ligand-binding domain of *NvGABAAR1*. This F0 male was crossed to a homozygous F3 *NvSH3GL3^P^>mScarlet-I* reporter female, and genotyping of 3–10 dpf F1 offspring confirmed germline transmission of the +27 allele. The remaining F1 siblings were raised to juvenile stages and individually genotyped to identify heterozygous *NvGABAAR1*^+/+27^, reporter-positive individuals. All such F1 animals were pooled and intercrossed to generate the F2 generation.

Individual F2 animals were genotyped to distinguish reporter-positive WT (*NvGABAAR1*^+/+^), heterozygous (*NvGABAAR1*^+/+27^), and homozygous (*NvGABAAR1*^+27/+27^) siblings. F3 WT controls were generated by intercrossing reporter-positive F2 WT siblings (*NvGABAAR1*^+/+^ × *NvGABAAR1*^+/+^), and F3 homozygous mutants were generated by intercrossing reporter-positive F2 homozygous siblings (*NvGABAAR1*^+27/+27^ × *NvGABAAR1*^+27/+27^). All neural-morphology imaging and behavioral assays were conducted on F3 homozygous mutants together with their F3 WT siblings. Throughout, “wild-type” refers to same-generation siblings lacking the +27 allele rather than unrelated WT animals.

For genotyping, whole embryos (<10 dpf) or aboral tissue from juveniles and adults was lysed in QuickExtract DNA Extraction Solution (Biosearch Technologies, QE09050) at 65 °C for 30 min and 98 °C for 2 min. The *NvGABAAR1* target locus was amplified by PCR, followed by a second round of amplification to introduce sample-specific dual barcodes. Amplicons were pooled and size-selected using the ProNex Size-Selective Purification System (Promega), quantified on a Qubit Fluorometer (Thermo Fisher Scientific), and assessed for size and purity on an Agilent Bioanalyzer. Purified pools were sequenced on an Illumina MiSeq 2 × 250 bp flow cell. The resulting sequence data were demultiplexed, read pairs were joined, and indel frequencies and allele identities were analyzed using CRIS.py^54^.

All primers used for sgRNA template generation and genotyping are listed in Supplementary Table 9.

### Behavioral assays

#### Animal preparation

Adult F3 *NvGABAAR1*^+27/+27^ mutants and their WT siblings were used for all behavioral experiments. Animals were ∼1 year post-fertilization and of comparable size; unusually large or small individuals were excluded. Because *Nematostella* lacks reliable external sex markers and no sex-specific differences in feeding behavior are known, mixed-sex animals were used.

Mutant and WT groups were maintained separately at RT in Pyrex glass dishes under low ambient light, as described above, and kept on the same shelf to minimize environmental variation. Animals were fed freshly hatched *Artemia* once per week. To avoid physiological effects of high-protein or non-native food sources (such as egg white), feeding assays were spaced by at least 4 weeks of routine *Artemia* feeding.

For each experiment, individual animals were transferred 2–4 days before the assay into sterile 60 × 15 mm polystyrene dishes (Falcon 351007) containing 12 ppt ASW and allowed to acclimate to the brighter ambient illumination near the imaging setup. Seawater was replaced the night before or morning of the assay to remove gametes and metabolic waste. On the day of testing, dishes were placed next to the imaging setup for ≥ 1 h so animals could fully relax and extend under the recording illumination, which was essential to prevent retraction during dish handling and the assay.

Animals were assayed 7, 11, or 14 days after their last *Artemia* feeding, with 14-day animals representing the more strongly starved condition. Mutant and WT animals from the same cohort were always tested in parallel on the same day, and datasets pooled measurements from multiple independent assay days.

#### Food preparation

For qualitative behavioral observations, *Artemia* nauplii were used for naturalistic feeding (Supplementary Video 1), whereas mussel tissue, torn into small pieces with fine forceps, was used for assays involving multiple food items, lateral presentation, and sequential feeding (Supplementary Videos 3–5).

For assays involving standardized food geometry, cylindrical egg-white pellets were prepared from the white portion of hard-boiled chicken eggs (Supplementary Videos 2 and 6). Egg-white slices (5–8 mm thick) were cored using an EMS Core Sampling Tool (EMS 69039) and gently pressed into a soft substrate to avoid compression artifacts, generating cylindrical logs. Logs were then cut into pellets with height equal to diameter using a 0.125-mm ultra-fine tungsten dissecting needle (Roboz Surgical RS-6063). Pellets of 0.5, 0.75, 1.0, 1.2, 1.5, and 2.0 mm diameter were used for size-series comparisons (Supplementary Video 2). A diameter of 1.0 mm was selected for all quantitative assays (Supplementary Video 6), as smaller pellets were difficult to track and did not reliably elicit mouth opening, whereas larger pellets introduced variability due to prolonged and inconsistent transit through the pharynx.

#### Food delivery

Food pellets were delivered by either a pipette- or whisker-based approach. In the pipette method, a pellet was released from a glass Pasteur pipette positioned adjacent to a tentacle near the oral opening. Glass pipettes were required because plastic pipettes caused egg-white pellets to adhere to the inner surface and impeded reliable release. This method produced a slight water disturbance, providing both mechanical and chemical cues. In the whisker method, the pellet was first released distally in the dish and then gently guided toward the oral opening using a trimmed cat whisker affixed to a wooden toothpick. The whisker provided sufficient rigidity for precise manipulation while introducing minimal mechanical stimulation. This method enabled more consistent placement of the food stimulus, although grasping onset was typically slower and more variable due to reduced mechanical cues relative to pipette delivery.

For swallowing analyses, pipette- and whisker-delivered trials were pooled because the delivery method did not significantly affect swallowing durations within each genotype. This was confirmed by the linear mixed-effects models (see Supplementary Table 8). Parallel to these findings, while delivery method influenced grasping durations, the effect was comparable across genotypes (Supplementary Fig. 5f).

#### Feeding assay

Dishes containing single acclimated animals were arranged beside the imaging setup and moved into view with minimal disturbance. All assays were recorded. Food pellets were delivered by either the pipette or whisker method, as described above. After a pellet was grasped and swallowed, the next trial began only once the ingested pellet had fully passed through the pharynx into the gastrovascular cavity and no further downward movement was observed.

A trial was defined as one pellet presentation and the resulting grasping and swallowing sequence. Each behavioral session (one animal per session) consisted of 4–10 sequential trials, but only the first four consecutive trials (trials 1–4) were used for quantitative analyses. Grasping and swallowing latencies showed no systematic change after the first three trials, and additional trials increased the likelihood of movement artifacts or handling disturbances.

Sessions were excluded if swallowing was accompanied by pronounced bending of the oral region, which mechanically altered ingestion dynamics. Such bending was typically triggered when a centrally delivered pellet was captured by a peripheral tentacle. To avoid these artifacts, any peripheral capture within trials 1–4 led to exclusion of the session. Peripheral capture occurring only after trial 4 did not affect inclusion, as later trials were not analyzed. Sessions were also terminated if the animal retracted into its body column during the assay, even if it later re-extended. Animals with unusually narrow pharynges were excluded to prevent anatomical constraints from influencing timing measurements.

#### Video acquisition

Behavioral sessions were recorded on a Leica M165 FC stereo microscope equipped with a Flexacam C1 camera. Videos were acquired at 1× magnification and 0.73× zoom, at either 0.50 or 0.87 s/frame.

#### Behavior annotation

Grasping and swallowing were quantified from time-lapse recordings as two sequential phases of the feeding response. For quantification, grasping was defined as the period from the first tentacle-driven engagement of the pellet to the onset of neck constriction, and swallowing as the period from the initiation of neck constriction to the moment the pellet reached the distal end of the pharynx.

For each trial, three frames were manually identified in Fiji:

t1 — first clear grasping movement after pellet introduction, marked by the onset of a tentacle-driven pull toward the mouth.

t2 — beginning of neck constriction, often coinciding with or following a tentacle clasping motion.

t3 — frame in which the pellet reached the distal end of the pharynx. This endpoint was used because the time from this point to entry into the gastrovascular cavity varied widely across trials and individuals.

Grasping and swallowing durations were calculated as the frame difference multiplied by the corresponding seconds-per-frame value: Tgrasp = (t2 − t1) × s/frame and Tswallow = (t3 − t2) × s/frame. Frame numbers and timestamps were recorded in Excel for downstream analysis. All trial-level grasping and swallowing duration measurements are provided in Supplementary Table 7.

#### Visualization and statistics

Data processing and visualization were performed in R (v4.4.1) using ggplot2 (v3.5.2) and standard tidyverse packages. Individual trial trajectories were plotted as faint lines, and genotype-wise mean ± s.e.m. were overlaid as solid lines. Grasping plots included pipette-delivery trials only, whereas swallowing plots included both pipette- and whisker-delivery methods.

All statistical analyses were performed in R. All statistical tests were two-sided. Feeding durations were analyzed using linear mixed-effects models fitted with lme4^55^ (v1.1-38). AnimalID was included as a random intercept to account for repeated measurements from the same individual across trials. Fixed effects included genotype and trial number. For swallowing analyses, food delivery method and starvation duration were included as additional covariates.

Significance of main effects and interactions was assessed using Type III Analysis of Variance (ANOVA) with Satterthwaite’s method implemented in lmerTest^56^ (v3.1-3). Genotype differences at each trial were estimated using estimated marginal means and pairwise contrasts computed with emmeans^57^ (v2.0.0), utilizing the Kenward-Roger approximation for degrees of freedom. The resulting estimates, standard errors, confidence intervals, and p-values are provided in Supplementary Table 8. Statistical inference relied on the assumptions of linear mixed-effects models; no additional tests of normality or global multiple-comparison corrections were performed.

### Whole-mount staining

Whole-mount staining was performed following previously published protocols^58^ for *Nematostella* with minor modifications. Planula larvae (3 dpf) and primary polyps (7–10 dpf) were collected into 1 ml ASW in round-bottom tubes. Primary polyps were allowed to relax for ≥ 15 min, after which MgCl_2_ was added stepwise by two sequential additions of 100 μl of 7% MgCl_2_ in 12 ppt ASW, with a 5-min incubation between additions. Planula larvae were immobilized by adding 7% MgCl_2_ directly, without a relaxation period. After immobilization, the solution was removed and replaced with fixative.

For visualization of F-actin in fixed transgenic reporter animals, samples were fixed in 4% paraformaldehyde (PFA) in 12 ppt ASW for 1 h at RT with gentle rocking, followed by washing 5 × 5 min in phosphate-buffered saline (PBS) containing 0.1% Tween-20 (PBTw). Samples were then incubated overnight at 4 °C on a shaker with either Alexa Fluor 488-conjugated phalloidin (1:300) or SiR-actin (Cytoskeleton, CY-SC001; 1:1000), washed again in PBTw, and cleared by immersion in ScaleA2 (2 M urea, 75% glycerol in water) prior to mounting and imaging.

For GABA immunostaining, animals were fixed for 3 min in 4% PFA supplemented with 0.05% glutaraldehyde in 12 ppt ASW, followed by post-fixation in 4% PFA in 12 ppt ASW for 1 h at room temperature with gentle rocking. Samples were washed 5 × 5 min in PBTw and incubated in blocking buffer (PBTw supplemented with 5% normal goat serum, 1% bovine serum albumin, and 10% dimethyl sulfoxide [DMSO]) for 2 h at RT. Primary antibody incubation was performed overnight at 4 °C on a shaker using rabbit anti-GABA (Sigma-Aldrich, A2052; 1:500) diluted in blocking buffer without DMSO. The following day, samples were washed 6 × 20 min in PBTw and incubated overnight at 4 °C on a shaker with goat anti-rabbit Alexa Fluor 647 secondary antibody (Invitrogen, A-21245; 1:1000) diluted in blocking buffer without DMSO. Samples were then washed 6 × 20 min in PBTw and cleared in ScaleA2 prior to imaging.

### Whole-mount fluorescent in situ hybridization

Gene-specific probe regions were selected and verified for specificity by BLAST searches against the *Nematostella* transcriptome. Nested PCR on cDNA templates amplified the target region, and the second-round reverse primer incorporated a T7 promoter sequence for in vitro transcription. Final PCR products were Sanger-sequenced to confirm identity. DIG-labeled antisense RNA probes were generated by in vitro transcription using the DIG RNA Labeling Mix (Roche, 11277073910), followed by DNase I treatment and column purification (Zymo Research, R2052). Primer sequences used for probe generation are listed in Supplementary Table 9.

FISH was performed based on established *Nematostella* protocols^59^. In brief, animals were immobilized as described for immunostaining, pre-fixed for 90 s in ice-cold 4% PFA + 2.5% glutaraldehyde in 12 ppt ASW, and then fixed in 4% PFA in 12 ppt ASW for 1 h at RT. Samples were washed in PBS and dehydrated through a PBS/methanol series into 100% methanol. Endogenous peroxidase activity was quenched in 3% H_2_O_2_ in 90% methanol for 1 h at RT, followed by rehydration through a methanol/PBS series. Tissues were permeabilized with Proteinase K (20 μg ml^-1^ for 10 min), post-fixed in 4% PFA for 1 h, washed in PBTw, and transitioned into hybridization buffer at 60 °C, where they were pre-hybridized overnight. DIG-labeled probes (1 ng μl^-1^) were denatured at 80 °C for 10 min, snap-cooled on ice, and hybridized to samples at 60 °C for 48 h. After hybridization, samples were washed at 60 °C through SSC solutions of increasing stringency, then returned to RT and washed into TNT (0.1 M Tris-HCl pH 7.5, 0.15 M NaCl, 0.1% Tween-20). Samples were blocked in TNT containing 5% sheep serum and 1% blocking reagent (Roche) for 2 h at RT, then incubated overnight at 4 °C with anti-DIG-POD Fab fragments (Roche, 11207733910; 1:1000). Fluorescent signal was developed using the TSA Cyanine 3 system (Akoya, NEL704A001KT) for 20 min at RT. After washing into PBTw, samples were counterstained with SiR-DNA (Cytoskeleton, CY-SC007; 1:1000), cleared in ScaleA2, and mounted for imaging as described for immunostaining.

### Imaging and image processing

All samples were imaged on either a Nikon spinning-disk confocal system (20× or 40× objectives) or an Andor Dragonfly spinning-disk confocal microscope (25× or 40× objectives). Z-stacks were acquired at 1-μm step size. Maximum intensity projections, orthogonal views generated by reslicing, and all subsequent image processing (rotation, cropping, and linear min/max intensity scaling) were performed in Fiji for visualization.

### Single-cell RNA-seq

#### Tissue dissociation and cell sorting

Adult F2 *NvSH3GL3^P^>mScarlet-I* animals were used for the single-cell RNA-seq experiment. All animals were of mixed sex, and the workflow was performed as a single batch on the same day. Animals were fasted for five days and transferred from 16 °C to RT for ∼8 h before dissociation. Animals were kept in 12 ppt ASW until fully relaxed and immobilized by gradually adding a 7% MgCl_2_ solution prepared in ASW (Sigma-Aldrich, M2670). Oral discs were dissected by a single transverse cut just below the pharynx, with most tentacles trimmed away. Dissected tissues were briefly rinsed in Ca^2+^/Mg^2+^-free ASW (154 mM NaCl, 3.57 mM KCl, 0.70 mM NaHCO_3_, 2.33 mM Na_2_SO_4_; all salts from Sigma-Aldrich), then briefly ground (∼15 s) and gently triturated for ∼20 min at RT in Ca^2+^/Mg^2+^-free ASW containing L-cysteine (2 mg ml^-1^; Sigma-Aldrich, 168149), papain (4 mg ml^-1^; Sigma-Aldrich, P4762), and Liberase TM (0.05 mg ml^-1^; Roche, 5401119001). Cell suspensions were filtered through 30 µm filters (FILCONS 030-33s; CTSV), pelleted (300 × g, 10 min, 4 °C), washed in enzyme-free Ca^2+^/Mg^2+^-free ASW, and stained on ice for 5 min with DAPI (2 µg ml^-1^; Sigma-Aldrich, D9542) and DRAQ5 (25 µM; Thermo Fisher Scientific, 62251).

Cells were analyzed and sorted on a BD Influx flow cytometer (100 µm nozzle, 20 psi). Excitation and emission were collected for DAPI (345/455 nm), DRAQ5 (647/697 nm), and mScarlet-I (578/613 nm). Because *Nematostella* oral tissue exhibits strong autofluorescence from the endogenous red fluorescent protein NvFP-7R^37^ (569/593 nm), gating was anchored using a fluorescence-minus-one (FMO) control prepared from WT oral-disc cells stained with DAPI and DRAQ5 but lacking mScarlet-I. This control captured the autofluorescence range of NvFP-7R and defined the background threshold for the reporter channel. A tentacle sample expressing mScarlet-I but lacking NvFP-7R established the mScarlet-I emission profile. Reporter-positive gates retained events bright in the mScarlet-I detector and dim in an adjacent 670/30 channel, minimizing NvFP-7R carryover. Only singlets and live, nucleated events (DAPI^-^/DRAQ5^+^) were included. The final gate contained ∼2% of total events in the reporter population versus 0.1% in the FMO control, yielding approximately 45,700 sorted events collected into RNase-free tubes for single-cell RNA-seq.

#### Library preparation and sequencing

Dissociated and sorted cells were assessed for concentration and viability using a LUNA-FL Cell Counter (Logos Biosystems). Samples with ≥ 90% viable cells were loaded onto a Chromium Controller (10x Genomics), targeting recovery of ∼3,200 cells. Single-cell cDNA libraries were prepared with the Chromium Single Cell 3′ Reagent Kit v3 (10x Genomics) according to the manufacturer’s instructions. cDNA and final short-fragment libraries were evaluated for quality and quantity using a 2100 Bioanalyzer (Agilent Technologies) and a Qubit Fluorometer (Thermo Fisher Scientific). Libraries were sequenced on an Illumina NextSeq 500 to a mean depth of ≥ 100,000 reads per cell (paired-end 28 × 8 × 91 bp) using Real-Time Analysis (RTA) and instrument software versions current at the time of sequencing.

FASTQ files were processed with Cell Ranger (v8.0.1; 10x Genomics, https://www.10xgenomics.com/) against the *Nematostella vectensis* genome assembly jaNemVect1.1 (GCF_932526225.1; NCBI RefSeq^60^ Annotation Release 101), supplemented with the *NvSH3GL3^P^>mScarlet-I* reporter sequence. The resulting gene–cell count matrix was analyzed in Seurat^61^ (v5.1.0) in R (v4.4.1). Cells were retained if they expressed 200–3,000 genes, contained ≤ 15,000 unique molecular identifiers (UMIs), and exhibited < 5% mitochondrial gene content (identified from RefSeq-annotated mitochondrial gene prefixes). Data were normalized with SCTransform^62^, and the top 3,000 most variable genes were used for principal-component analysis (PCA).

The first 15 principal components were used to construct a shared nearest-neighbor graph (FindNeighbors) and perform graph-based clustering (FindClusters, resolution = 0.5). Uniform Manifold Approximation and Projection (UMAP) embeddings computed from the same components were used for visualization.

#### Clustering and analysis

Cluster identities were assigned by examining marker genes identified with FindAllMarkers and cross-referenced with manually curated *Nematostella* gene sets for neuronal developmental, synaptic function, sensory transduction, and neurotransmitter receptors (Supplementary Tables 1 and 2). These curated lists were generated from NCBI RefSeq annotations using keyword-based filtering and provided the framework for assigning biologically meaningful identities to each cluster.

Gene-set quantification was performed using the same manually curated gene lists. Expression was evaluated from the SCT-normalized count layer, and a gene was considered detected in a cluster if it exhibited non-zero expression and was expressed in more than 10% of cells in that cluster. For each gene set, per-cluster counts of detected genes and per-gene summary tables (average expression, percent of expressing cells, and annotation metadata) are provided in Supplementary Table 2.

All visualizations were generated in R using Seurat (v5.1.0), ggplot2 (v3.5.2), ggtern (v3.5.0), pheatmap (v1.0.12), patchwork (v1.3.0), and standard tidyverse packages. All analysis and visualization code is available at [link].

### Bulk RNA-seq

Adult male *Nematostella vectensis* polyps (n = 3 individuals) were dissected into four tissue types (pharynx, body wall, proximal tentacles, and oral disc), yielding twelve total samples (three biological replicates per tissue). Each tissue piece was immersed in 300 µl TRIzol reagent (Invitrogen, 15596026) and snap-frozen in liquid nitrogen before storage at –80 °C. RNA was extracted using the Direct-zol RNA MiniPrep Kit (Zymo Research, R2052) following the manufacturer’s protocol, including a 15 min on-column DNase I digestion to remove genomic DNA.

mRNA-seq libraries were generated from 100 ng (or < 100 ng for two samples) of high-quality total RNA, as assessed using the Bioanalyzer (Agilent). Libraries were prepared using the TruSeq Stranded mRNA Library Prep Kit (Illumina, 20020594) and TruSeq RNA Single Indexes Sets A and B (Illumina, 20020492 and 20020493). Libraries were purified using the AMPure XP bead-based reagent (Beckman Coulter, A63882), evaluated for quality and quantity using the Bioanalyzer (Agilent) and Qubit Fluorometer (Life Technologies). Equal-molar libraries were pooled, quantified, and sequenced as 75 bp paired-end reads on a mid-output flow cell using the NextSeq 500 (Illumina). Following sequencing, demultiplexing and FASTQ generation were performed using Illumina RTA (v2.4.11) and bcl2fastq2 (v2.20).

FASTQ files were aligned to the *Nematostella vectensis* genome assembly jaNemVect1.1 (GCF_932526225.1; NCBI RefSeq^60^ Annotation Release 101) using STAR^63^ (v2.7.10b) with quantMode GeneCounts to obtain gene-level counts. Alignment summary rows were removed and raw counts were imported into R and analyzed with edgeR^64^ (v4.2.2). Lowly expressed genes were filtered using filterByExpr with tissue as the grouping factor, and library sizes were normalized using the trimmed mean of M-values (TMM) method. Gene lengths were obtained from the corresponding RefSeq gene models and used to compute TPM values from the filtered count matrix (Supplementary Table 3). For visualization, TPM values were averaged across biological replicates for each tissue, restricted to a curated list of GABA receptor-like subunits (Supplementary Table 4), and scaled per gene (row-wise) before generating clustered heatmaps.

### Phylogenetic analysis

#### Sequence identification and curation

Putative *Nematostella vectensis* γ-aminobutyric acid type A receptor (GABA_A_R) and glycine receptor (GlyR) subunits were first identified by BLASTp^65^ searches (v2.13.0+) against the *Homo sapiens* GABA_A_R and GlyR sequences and verified for domain composition using the InterPro server^66^. Although orthology between GABA_A_R and GlyR types remains unresolved in early-branching metazoans, no hits overlapped with nicotinic acetylcholine, 5-HT_3_, or other members of the Cys-loop ligand-gated ion channel (LGIC) superfamily. These sequences are therefore collectively referred to as GABA_A_R-like subunits throughout.

For *Nematostella*, candidate GABA_A_R-like sequences were compiled from three curated sources: the NCBI RefSeq proteome^60,19^ (https://www.ncbi.nlm.nih.gov/datasets/genome/GCF_932526225.1/) and two SIMRBase transcriptome-derived proteome assemblies^18^ (NV2: wein_nvec200_tcsv2; NVE: wien_nvec200_v1; https://simrbase.stowers.org/starletseaanemone). Not all loci were represented across all datasets. When the same locus appeared in multiple sources, we retained the longest predicted amino-acid product; when sequences were equal in length, the RefSeq entry was selected. This procedure yielded 40 curated *Nematostella* GABA_A_R-like subunits: 37 for which the RefSeq version was retained, 2 for which RefSeq lacked annotated protein entries and the longest ORFs were contributed by the NV2 assembly, and 1 for which the longest ORF was contributed by the NVE assembly, despite orthologous predictions also present in RefSeq and NV2 (Supplementary Table 4).

Homologous GABA_A_R-like sequences for comparative phylogenetic analysis were retrieved from UniProt^67^ (Swiss-Prot and TrEMBL; downloaded February 2024) for six bilaterian species (*Homo sapiens*, *Danio rerio*, *Drosophila melanogaster*, *Caenorhabditis elegans*, *Ciona intestinalis*, and *Octopus vulgaris*). The *Nematostella* and bilaterian sequences were used as queries for BLASTp^65^ searches against six additional cnidarian proteomes representing both anthozoans (*Acropora millepora*, *Exaiptasia diaphana*, *Stylophora pistillata*) and medusozoans (*Clytia hemisphaerica, Hydra vulgaris*, *Hydractinia symbiolongicarpus*). Resulting sequences from these searches were further curated using CD-HIT^68,69^ (v4.8.1) to remove redundant sequences sharing ≥ 90 % similarity within each species. Protein database sources, accession numbers, and annotations for all taxa are listed in Supplementary Table 5.

#### Sequence alignment and phylogeny reconstruction

The non-redundant candidates from all species were aligned with MAFFT^70^ (v7.520) using the L-INS-i algorithm, and discordant sequences were excluded after manual inspection. Domain composition was re-evaluated using the InterPro server^66^ (https://www.ebi.ac.uk/interpro/search/sequence/, accessed 19 Nov 2024), and sequences matching the GABA_A_R/GlyR α-subunit domain (IPR006028) were retained. No matches to the GlyR β-subunit domain (IPR008060) were detected. The final alignment was trimmed with TrimAl^71^ (v1.4.rev22) using the -gappyout flag.

A maximum-likelihood phylogeny was constructed using the IQ-TREE 2 server^72^ (v1.6.12; http://iqtree.cibiv.univie.ac.at/, accessed March 2025) under the LG+I+G4 substitution model selected by ModelFinder^73^. Branch support values were estimated from 1,000 replicates using ultrafast bootstrap (-bb)^74^ and SH-like approximate likelihood-ratio tests (-alrt)^75^. Representative nicotinic acetylcholine receptor (nAChR) subunits were included as an outgroup to root the tree. The resulting phylogeny was visualized in FigTree (v1.4.4; http://tree.bio.ed.ac.uk/software/figtree/) and formatted using Inkscape (v1.3; www.inkscape.org) and Adobe Illustrator (v30.0).

### AlphaFold structure prediction

The predicted structure of the *Nematostella vectensis* GABA_A_R-like subunit NvGABAAR1 was generated using the AlphaFold3 web server^76^ (https://alphafoldserver.com, accessed June 2025), using the longest annotated RefSeq isoform (gene ID: LOC5505184; protein ID: XP_048587017.1) as input. Predictions were run with default parameters, and the top-ranked structural model was used for visualization and structural comparison.

For cross-species comparison, AlphaFold-predicted structures of *Homo sapiens* GABA type A receptor subunits GABRR3 (UniProt A8MPY1), GABRA2 (UniProt P47869), and GABRG1 (UniProt Q8N1C3) were obtained from the AlphaFold Protein Structure Database in July 2025. All human structures correspond to AlphaFold models version v4 (AF-*-F1-v4).

Structural visualization and pairwise alignment were performed in UCSF ChimeraX^77^ (v1.9). All models were colored by per-residue model confidence, measured as predicted Local Distance Difference Test (pLDDT), and aligned using the MatchMaker algorithm.

## Use of AI tools

Large language models (ChatGPT, OpenAI) were used under author direction to assist with code generation, debugging, and improving the clarity of written text. All scientific analyses, code execution, and validation were performed by the authors.

## Data availability

Single-cell RNA-seq and bulk RNA-seq datasets generated in this study have been deposited in the Gene Expression Omnibus (GEO) under accession numbers GSE316974 and GSE316975. All source data will be deposited in the Stowers Original Data Repository (ODR) prior to publication. Additional supporting data are available from the corresponding author upon request.

## Code availability

Custom R scripts used for data analysis and figure generation are provided with this submission and will be made publicly available upon publication.

## Acknowledgements

We thank Boris Rubinstein and Kausik Si for discussions on behavioral experiments, and Kausik Si additionally for advice on neurotransmitter receptor composition analysis. We thank Joshua Zimmerberg for discussions on body size and nerve net scalability, and Neşet Özel for insights into the relevance of the vesicular GABA transporter in shaping interneuron identity. We are grateful to Mark Miller for creating the *Nematostella* life-cycle illustration, and to past and present members of the Gibson lab for discussions and support. We acknowledge the Stowers Technology Centers for assistance: the Aquatics team for animal husbandry; the Cytometry team, particularly Jillian Blanck, for cell sorting; the Sequencing and Discovery Genomics team for library preparation and sequencing; the Computational Biology team for processing of sequencing data; the Genome Engineering team for CRISPR genotyping; and the Microscopy Center for imaging training and support. This work was supported by the Stowers Institute for Medical Research.

## Author contributions

R.Z. conceived and designed the study; performed all experiments and data analyses; prepared the figures; and wrote the manuscript. M.C.G. supervised the research, provided guidance, and edited the manuscript. C.W.S. performed preprocessing of sequencing data, generated genome-assembly mapping tables, and supervised code implementation. A.M.L.K. performed sequence alignment and phylogenetic tree construction. All authors reviewed and approved the final manuscript.

