## Supplementary Information for "Morphological, molecular, and functional evidence for a CNS-like oral nerve ring in the sea anemone *Nematostella vectensis*"

This PDF contains supplementary figures with their corresponding legends, descriptions of supplementary tables and videos, and supplementary notes.

**Supplementary Figures 1–5**
**Supplementary Tables 1–9**
**Supplementary Videos 1–6**
**Supplementary Notes**

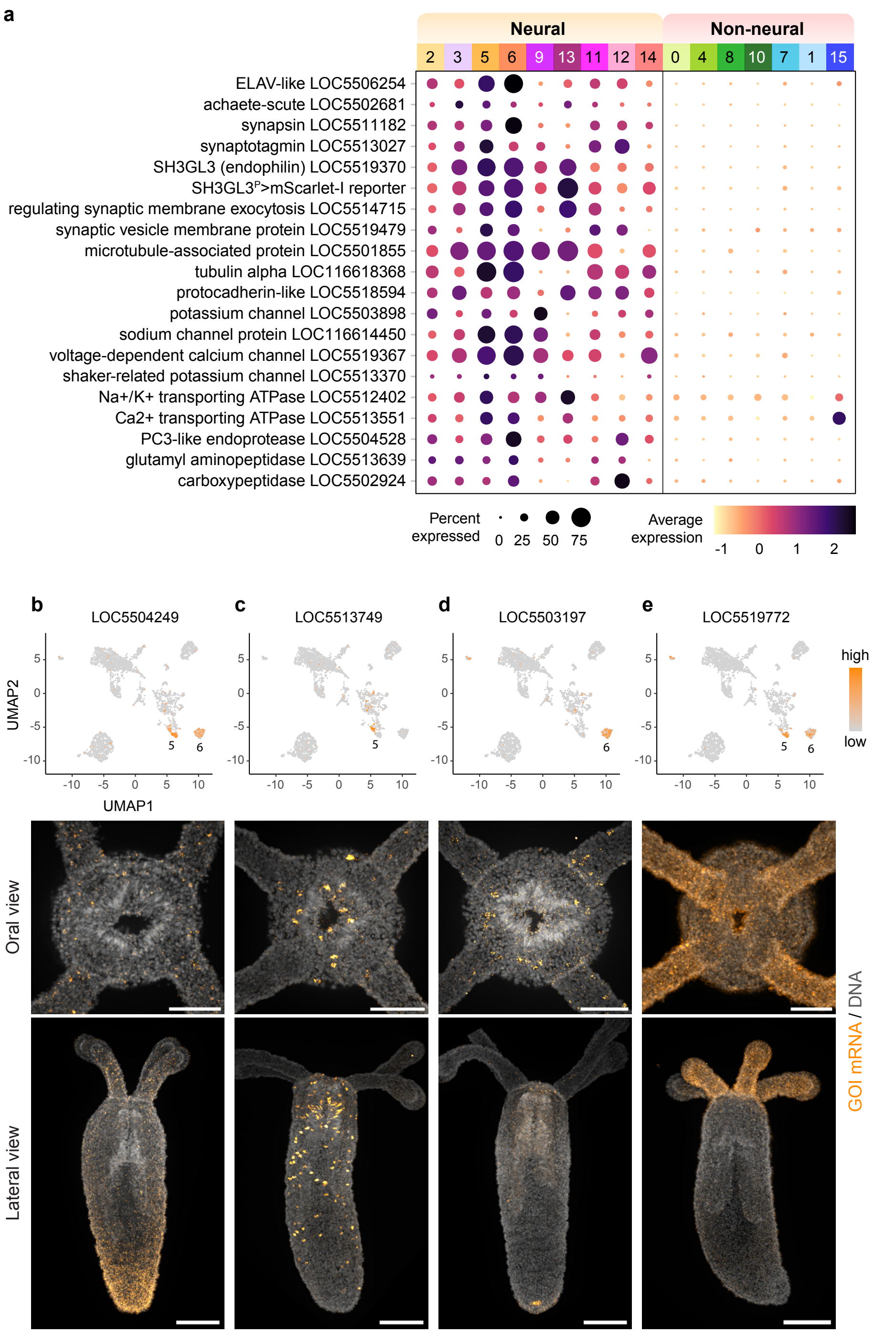


**Supplementary Fig. 1 | Molecular and spatial validation of oral neuronal cluster identities.**

**(a)** Dot plot showing expression of canonical neuronal markers, synaptic machinery, ion channels, cytoskeletal components, and signaling enzymes across neural and non-neural clusters. Dot size indicates the percentage of cells expressing each gene, and color denotes scaled average expression. **(b–e)** Representative fluorescent in situ hybridization (FISH) analyses in four-tentacle primary polyps (7–10 days post-fertilization) for genes enriched in clusters 5 and 6 that did not meet criteria for oral nerve ring (ONR) markers. Shown are examples of transcripts that are weak or undetectable at this stage (b), expressed in the body column (c,d), or diffusely distributed across oral tissues without enrichment in ONR-like neurons (e). Oral (top) and lateral (bottom) views are shown for each FISH experiment. DNA is shown in gray and gene-specific signal in orange. Scale bars, 50 μm (oral views) and 100 μm (lateral views).

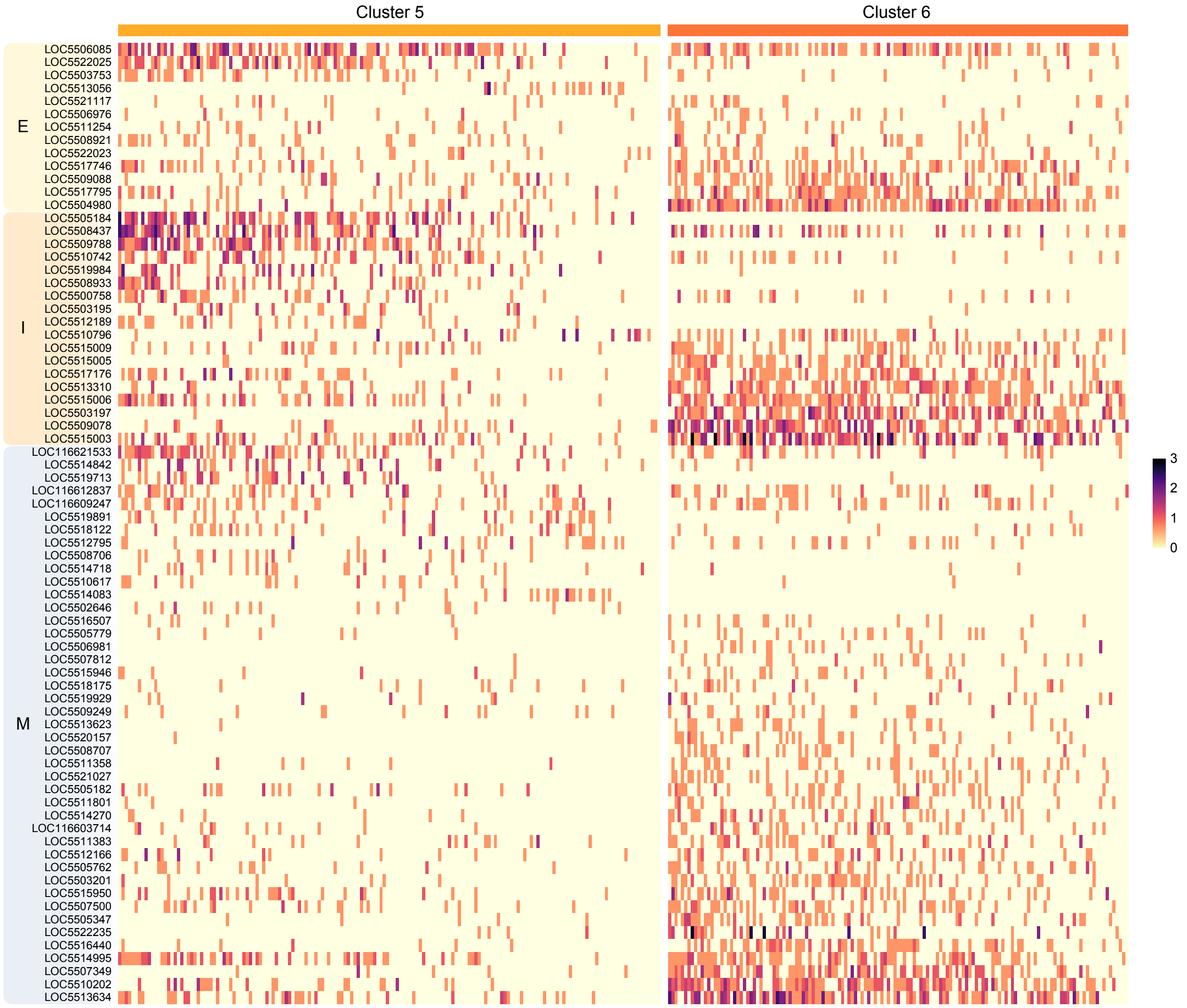


**Supplementary Fig. 2 | Neurotransmitter receptor family expression across interneuron-like clusters.**

Heatmap showing SCT-normalized expression of excitatory (E), inhibitory (I), and modulatory (M) neurotransmitter receptor genes across individual cells from clusters 5 and 6. Columns represent single cells, ordered within each cluster by aggregate expression of the displayed receptor set. Rows represent receptor genes, grouped by gene category and arranged from cluster 5-enriched (top) to cluster 6-enriched (bottom). Color indicates SCT-normalized expression (see color bar). This analysis provides gene-level context for the integrative receptor repertoires summarized in Fig. 4a–c and for the inhibitory signaling components examined in Fig. 6a.

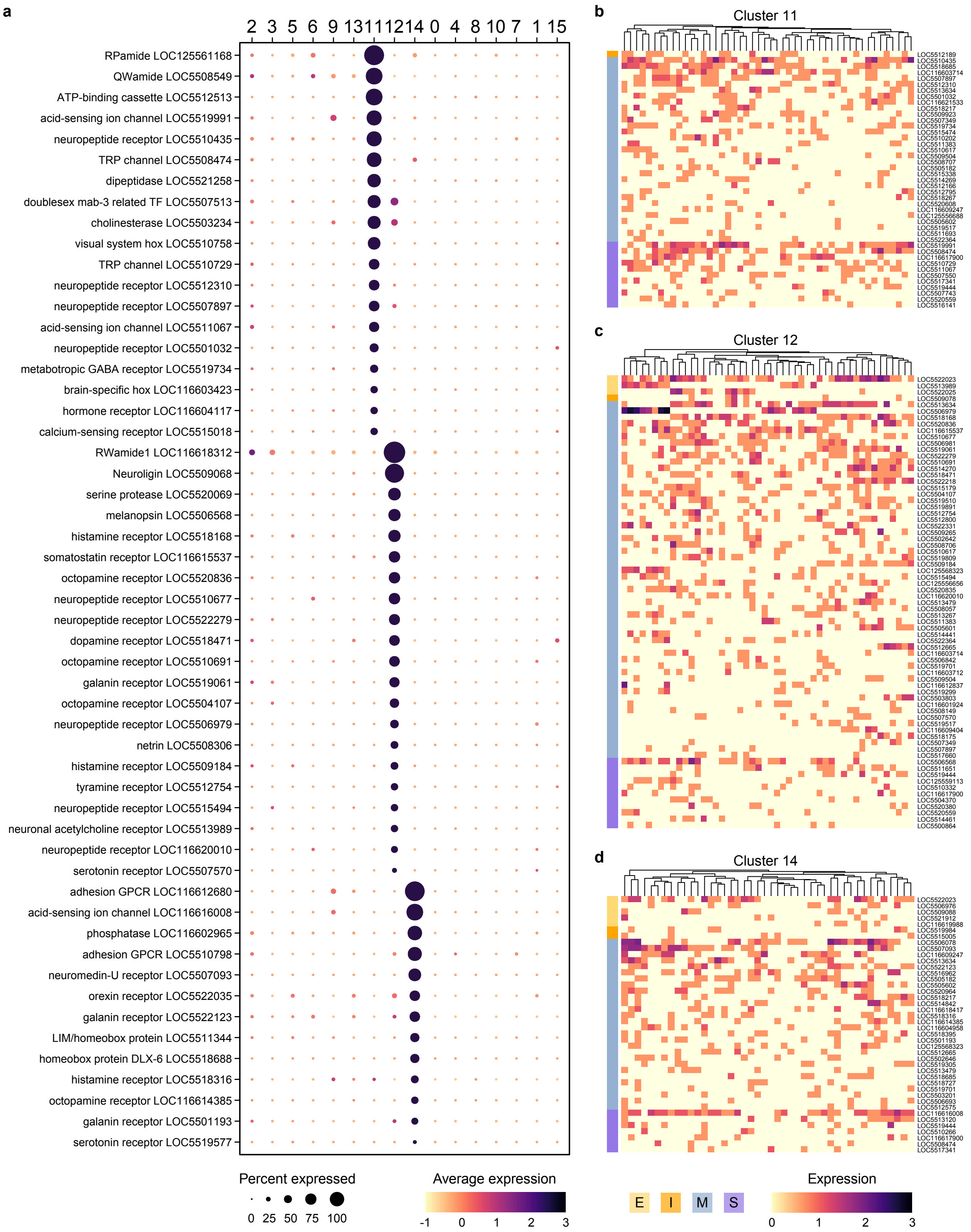


**Supplementary Fig. 3 | Molecular features of modulatory and sensory-enriched neuronal clusters.**

**(a)** Dot plot showing expression of marker genes enriched in clusters 11, 12, and 14. Dot size indicates the percentage of cells expressing each gene, and color represents scaled average expression (pale yellow to dark purple). **(b–d)** Heatmaps showing SCT-normalized expression of cluster-enriched genes in clusters 11 (b), 12 (c), and 14 (d). Columns represent individual cells ordered by hierarchical clustering within each cluster, and rows represent genes grouped by functional category. Gene groups are color-coded as ionotropic excitatory (gold), ionotropic inhibitory (orange), metabotropic/modulatory (blue), and sensory (purple).

51

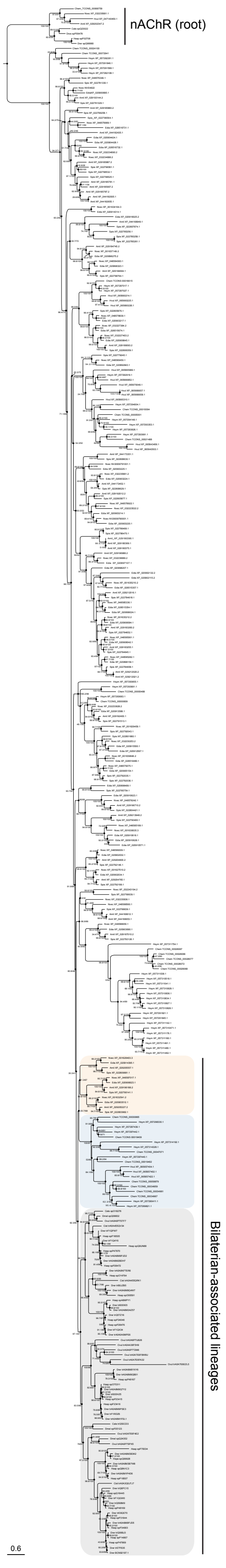

**Supplementary Fig. 4 | Full maximum-likelihood phylogeny of $GABA_A$R-like subunits.**

Maximum-likelihood reconstruction of bilaterian and cnidarian $GABA_A$R-like homologs inferred using IQ-TREE. Branch support at each node is shown as ultrafast bootstrap and SH-like aLRT values. Bilaterian-associated lineages are highlighted. Rounded background shading indicates major taxonomic groups within these lineages: grey shading indicates bilaterian branches, orange shading indicates anthozoan branches, and blue shading indicates medusozoan branches. Anthozoan and medusozoan branches outside bilaterian-associated lineages, comprising independently expanded cnidarian lineages, are shown without shading. Representative nicotinic acetylcholine receptor (nAChR) subunits were included as an outgroup to root the tree. This tree corresponds to the summary tree shown in Fig. 5e. Species abbreviations: Amil = *Acropora millepora*; Cele = *Caenorhabditis elegans*; Chem = *Clytia hemisphaerica*; Cint = *Ciona intestinalis*; Dmel = *Drosophila melanogaster*; Drer = *Danio rerio*; Edia = *Exaptasia diaphana*; Hsap = *Homo sapiens*; Hsym = *Hydractinia symbiolongicarpus*; Hvul = *Hydra vulgaris*; Nvec = *Nematostella vectensis*; Ovul = *Octopus vulgaris*; Spis = *Stylophora pistillata*.

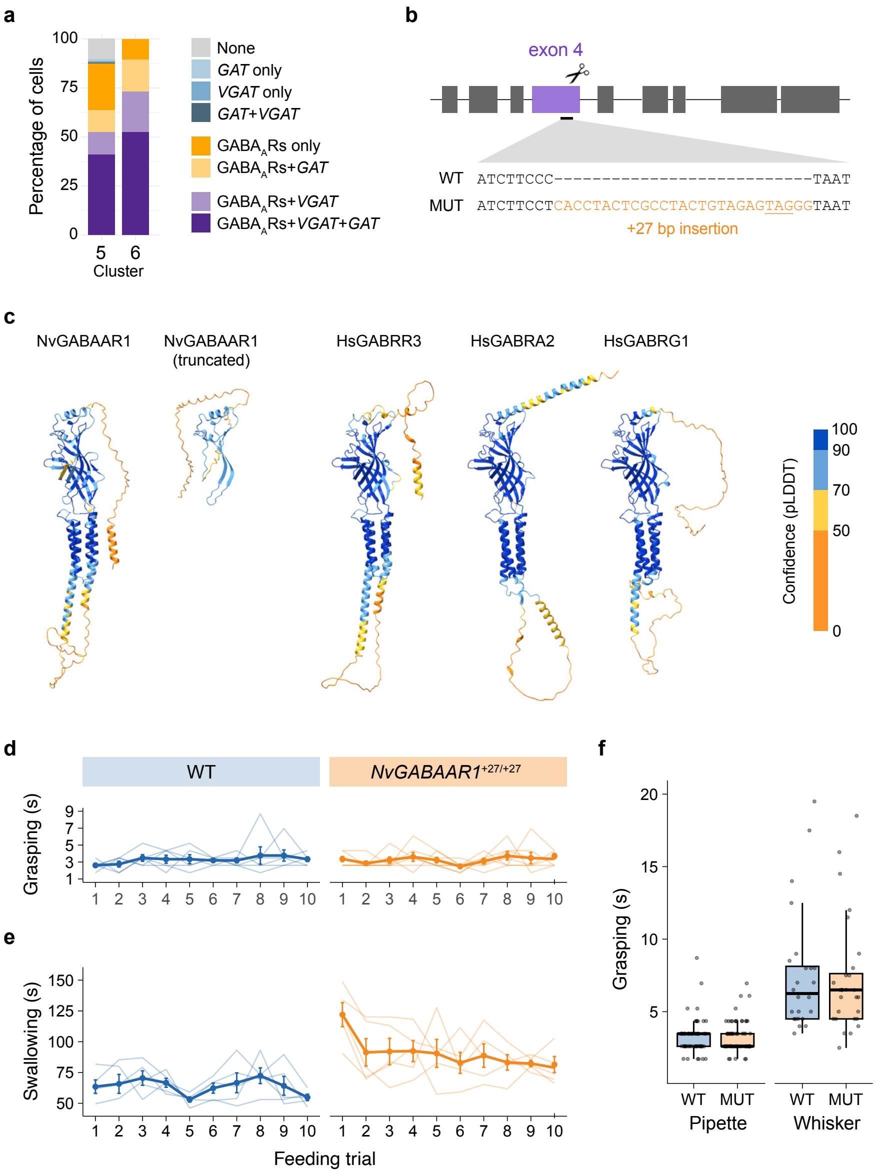


**Supplementary Fig. 5 | Molecular context, mutant design, and extended analysis of *NvGABAAR1*$^{+27/+27}$ feeding behavior.**

**(a)** Stacked bar plot showing the percentage of cells in clusters 5 and 6 expressing combinations of γ-aminobutyric acid (GABA) pathway components, including GABA type A receptor-like (GABA$_A$R-like) subunits, the vesicular GABA transporter (*VGAT*), and the plasma membrane GABA transporter (*GAT*). This analysis complements the single-cell-level expression patterns shown in Fig. 6a. “None” indicates cells lacking detectable expression of all three components. For brevity, GABA$_A$Rs in the color key refer to GABA$_A$R-like subunits. Colors denote distinct co-expression categories: none (gray); *GAT* only (light blue); *VGAT* only (blue); *GAT* + *VGAT* (dark blue); GABA$_A$Rs only (orange); $GABA_A$Rs + *GAT* (light orange); $GABA_A$Rs + *VGAT* (lavender); and $GABA_A$Rs + *VGAT* + *GAT* (purple). **(b)** Schematic of the *NvGABAAR1*$^{+27}$ allele. CRISPR/Cas9 targeting in exon 4 introduced a 27-bp insertion (orange), resulting in a frameshift and a premature stop codon (underlined). **(c)** AlphaFold-predicted structures of the full-length *Nematostella vectensis* $GABA_A$ receptor-like subunit NvGABAAR1, the truncated NvGABAAR1 protein generated by the +27-bp frameshift mutation shown in (b), together with representative human $GABA_A$ receptor subunits (HsGABRR3, HsGABRA2, and HsGABRG1), extending the structural comparisons shown in Fig. 5f. Structures are displayed as ribbon models and colored by per-residue model confidence (pLDDT), from low (orange) to high (blue). This comparison highlights the structural consequence of NvGABAAR1 truncation and illustrates the range of structural diversity among human $GABA_A$ receptor subunits. **(d)** Grasping duration across ten sequential feeding trials per animal in wild-type (WT; blue, n = 6) and *NvGABAAR1*$^{+27/+27}$ mutants (orange, n = 7), using pipette-delivered food pellets. Each thin line represents an individual animal; thick lines show the mean ± s.e.m. **(e)** Swallowing duration across ten sequential feeding trials per animal in wild-type (n = 5) and *NvGABAAR1*$^{+27/+27}$ mutants (n = 5), using pipette-delivered food pellets. Each thin line represents an individual animal; thick lines show the mean ± s.e.m. **(f)** Grasping duration for WT (blue) and *NvGABAAR1*$^{+27/+27}$ mutants (orange) under two pellet-delivery conditions. Pipette: WT (n = 6 animals, 60 trials) and mutants (n = 7 animals, 70 trials). Whisker: WT (n = 4 animals, 24 trials) and mutants (n = 5 animals, 28 trials). Each point represents an individual trial. Boxplots show the median (horizontal line) and interquartile range (IQR; 25th–75th percentile), with whiskers extending to the most extreme data points within 1.5× the IQR. All data points are plotted as horizontally jittered points.

**Supplementary Table 1**
scRNA-seq cluster markers and curated gene lists.
Related to Fig. 3, Fig. 4d, Supplementary Fig. 1, and Supplementary Fig. 3a.

**Supplementary Table 2**
scRNA-seq neurotransmitter receptor and sensory gene-set analyses.
Related to Fig. 4a–c, Supplementary Fig. 2, and Supplementary Fig. 3b–d.

**Supplementary Table 3**
TPM values underlying bulk RNA-seq expression analyses.
Related to Fig. 5c.

**Supplementary Table 4**
GABA receptor subunits in *Nematostella vectensis* (type A + type B).

**Supplementary Table 5**
GABA type A receptor-like subunit sequence sources and metadata.
Related to Fig. 5d,e and Supplementary Fig. 4.

**Supplementary Table 6**
Candidate glutamate decarboxylase (GAD) and GAD-like genes in *Nematostella vectensis* and their expression in oral scRNA-seq.
Related to Supplementary Notes.

**Supplementary Table 7**
Grasping and swallowing duration measurements.
Related to Fig. 6e,f and Supplementary Fig. 5d–f.

**Supplementary Table 8**
Statistical analyses of grasping and swallowing behaviors underlying Fig. 6.

**Supplementary Table 9**
Oligonucleotides used for reporter line construction, in situ hybridization, CRISPR mutagenesis, and genotyping.

**Supplementary Video 1**

Adult *Nematostella vectensis* partially burrowed in gravel feeding on free-swimming *Artemia* nauplii (related to Fig. 1b).

**Supplementary Video 2**

Feeding behavior with increasing cylindrical egg-white food pellet sizes (0.5, 0.75, 1.0, 1.2, 1.5, and 2.0 mm).

**Supplementary Video 3**

When multiple mussel food particles are presented within a brief interval, they are grouped and compressed into a single bolus prior to swallowing.

**Supplementary Video 4**

Feeding behavior when mussel food is presented from lateral positions, accomplished through collective movements of multiple tentacles and bending of the neck.

**Supplementary Video 5**

When a new mussel food stimulus is presented during ongoing swallowing, tentacle grasping is delayed and resumes after the swallowing is completed.

**Supplementary Video 6**

A simplified feeding paradigm in which a 1 mm × 1 mm cylindrical egg-white pellet is delivered to the oral disc using a cat whisker, eliciting a consistent feeding sequence used for behavioral analyses across genotypes (characterized in Fig. 1d). Images were acquired at 0.5 s per frame, and the video is shown at 5 frames per second (2.5× real time).

### Supplementary Notes

#### Behavioral flexibility during feeding

To probe behavioral flexibility, we manipulated prey characteristics and delivery methods under controlled conditions. When presented with size-standardized mock prey (cylindrical egg-white pellets), polyps engaged progressively more tentacles as prey size increased from 0.5 to 2.0 mm, indicating that motor recruitment scales with prey size (Supplementary Video 2). When multiple small food particles were presented within a brief interval, animals grouped and compressed them into a single bolus prior to swallowing (Supplementary Video 3). Responsiveness was also shaped by mechanical context (Supplementary Fig. 5f): direct pipette delivery near a tentacle elicited faster and less variable responses than distal delivery guided by a cat whisker (median latency 3.48 s vs. 6.25 s; IQR 0.87 s vs. 3.63 s), revealing sensitivity to both mechanical and chemosensory cues. This phenomenon also occurred during routine husbandry, where mild agitation enhanced feeding in four-tentacle polyps exposed to crushed *Artemia nauplii*. Animals further adjusted to prey position: when food was presented from lateral positions (Supplementary Video 4), feeding was achieved through collective movements of multiple tentacles and bending of the neck. Finally, when a new food stimulus was introduced during ongoing swallowing, tentacle grasping was delayed and resumed only after completion of the swallowing phase (Supplementary Video 5). Collectively, these observations demonstrate a high degree of motor flexibility and spatiotemporal coordination in an organism traditionally considered to lack centralized neural control.

#### Functional uncertainty of GAB$_A$A receptor-like subunits in *Nematostella*

Although GABA$_A$R-like subunits are abundant in *Nematostella vectensis*, their physiological properties remain largely uncharacterized. A recent preprint (Ojha et al., 2025) testing seven subunits reported that three were gated by glutamate rather than GABA, four showed no responses to any ligands tested, and the remaining 33 subunits, including the paralog we mutated, have not been examined. Because these assays relied exclusively on homomeric assemblies, which may not reflect native heteromeric configurations, non-responsiveness is difficult to interpret. Moreover, even if some subunits are gated by GABA, the functional sign of their activity cannot be inferred: inhibitory versus excitatory effects depend on channel ion selectivity and intracellular chloride gradients, neither of which is known for *Nematostella* neurons. Together, these considerations highlight the need for future electrophysiological and ion-substitution experiments to establish ligand identity, ion selectivity, and net physiological effects of $GABA_A$R-like subunits in the oral nerve ring (ONR).

**Search for canonical and non-canonical GABA synthesis pathways**

Bulk mass spectrometry detected GABA in whole-animal extracts (unpublished data). Despite the abundance and conservation of GABA transporters and receptors, the enzymatic route underlying GABA synthesis in *Nematostella* remains unresolved. All candidate genes identified in the analyses below, their correspondence to previously published loci, and their expression status in the oral scRNA-seq dataset are summarized in Supplementary Table 6.

We first searched for canonical glutamate decarboxylase (GAD) homologs based on protein sequence similarity to human and bacterial GADs. This analysis identified three low-confidence candidates: LOC5515329 and LOC5515304, both predicted to encode aromatic L-amino-acid decarboxylases, and LOC5519959, predicted to encode sphingosine-1-phosphate lyase. BLASTp searches using human GAD recovered high-confidence orthologs with strong sequence identity or query coverage in medusozoan cnidarians (*Hydra vulgaris*, *Hydractinia symbiolongicarpus*, *Clytia hemisphaerica*), but not in anthozoans (*Nematostella vectensis*, *Acropora millepora*, *Exaiptasia diaphana*, *Stylophora pistillata*), suggesting loss or substantial divergence of bilaterian-like GAD in the anthozoan lineage.

Previously reported *Nematostella* GAD or GAD-like homologs (v1g70014, v1g224555; Oren et al., 2014) and (v1g208016, v1g70058, v1g60834, v1g60452; Levy et al., 2020) now map to two loci in the latest NCBI RefSeq genome assembly, LOC5512039 and LOC5512040, respectively. Both loci are predicted to encode L-tyrosine decarboxylase-like enzymes rather than canonical GADs.

To capture more divergent candidates, we next searched for proteins containing the pyridoxal-dependent decarboxylase domain (Pfam PF00282), which is shared by GAD and related enzymes. This expanded search recovered the same five candidates noted above, together with five additional weak hits. However, none showed enrichment in any neuronal cluster in the adult oral disc scRNA-seq dataset, nor did any co-express with *VGAT*, a defining feature of GABAergic neurons. Although some candidates were detectable in adult bulk RNA-seq from dissected oral tissues, none were enriched in tentacles, where GABA immunoreactivity is strongest, further reducing their plausibility as neural GAD-like enzymes.

Finally, we evaluated potential non-canonical GABA synthesis routes. These included putrescine-derived pathways involving diamine oxidase (DAO) and aldehyde dehydrogenase (Aldh1a1), as well as monoamine oxidase B (MAO-B). Given the functional parallels between GABA and glycine as inhibitory neurotransmitters, and the structural similarity of their ionotropic receptors, we also examined serine hydroxymethyltransferase (SHMT), the key enzyme in glycine biosynthesis. However, none of these homologs showed strong or selective expression in neuronal populations within our scRNA-seq dataset. Together, these observations suggest that GABA production in *Nematostella* may rely on highly divergent, lineage-specific, or as-yet-uncharacterized enzymatic mechanisms.
